# Single-Cell Atlas of Common Variable Immunodeficiency reveals germinal center-associated epigenetic dysregulation in B cell responses

**DOI:** 10.1101/2021.12.20.473453

**Authors:** Javier Rodríguez-Ubreva, Anna Arutyunyan, Marc Jan Bonder, Lucía Del Pino-Molina, Stephen J. Clark, Carlos de la Calle-Fabregat, Luz Garcia-Alonso, Louis-François Handfield, Laura Ciudad, Eduardo Andrés-León, Felix Krueger, Francesc Català-Moll, Virginia C. Rodríguez-Cortez, Krzysztof Polanski, Lira Mamanova, Stijn van Dongen, Vladimir Yu. Kiselev, María T. Martínez-Saavedra, Holger Heyn, Javier Martín, Klaus Warnatz, Eduardo López-Granados, Carlos Rodríguez-Gallego, Oliver Stegle, Gavin Kelsey, Roser Vento-Tormo, Esteban Ballestar

## Abstract

Common variable immunodeficiency (CVID), the most prevalent symptomatic primary immunodeficiency, is characterized by impaired terminal B-cell differentiation and defective antibody responses. Incomplete genetic penetrance and a wide range of phenotypic expressivity in CVID suggest the participation of additional pathogenic mechanisms. Monozygotic (MZ) twins discordant for CVID are uniquely valuable for studying the contribution of epigenetics to the disease. We used single-cell epigenomics and transcriptomics to create a cell census of naïve-to-memory B cell differentiation in a pair of CVID-discordant MZ twins. Our analysis identifies DNA methylation, chromatin accessibility and transcriptional defects in memory B cells that mirror defective cell-cell communication defects following activation. These findings were validated in a cohort of CVID patients and healthy donors. Our findings provide a comprehensive multi-omics map of alterations in naïve-to-memory B-cell transition in CVID and reveal links between the epigenome and immune cell cross-talk. Our resource, publicly available at the Human Cell Atlas, paves the way for future diagnosis and treatments of CVID patients.

## INTRODUCTION

Common variable immunodeficiency (CVID), the most frequent symptomatic primary immunodeficiency^1^, is represented by a heterogeneous group of patients with low serum immunoglobulin concentrations, defective specific antibody production and increased susceptibility to bacterial infections of the respiratory and gastrointestinal tracts^2^. A high proportion of CVID patients have a low frequency of memory B cells^3^. In fact, CVID is mainly considered to be a humoral immunodeficiency, although several immune cell types might be affected ^4–7^.

CVID is usually detected sporadically in patients with no family history of immunodeficiency. However, for about 10% of subjects, other first-degree relatives may either be hypogammaglobulinemic or present selective IgA deficiency^8^. The existence of a genetic component in CVID has been recognized for decades and next-generation sequencing has revealed a number of pathogenic genes^9^. However, CVID pathogenesis remains largely unexplained, given that only 20% of CVID cases can be accounted for by monogenic gene defects, and genetic mutations remain elusive in the majority of CVID cases. The absence of mutations for a high proportion of CVID patients together with the incomplete disease penetrance for those harboring mutations suggest the operation of additional, as yet undefined, mechanisms (e.g., polygenic, epigenetic and environmental contributors) that help determine the CVID clinical phenotype.

We recently identified DNA methylation alterations associated with CVID, using a pair of monozygotic (MZ) twins discordant for the disease and a cohort of patients and controls, providing a concrete proof-of-principle of an epigenetic dimension to CVID^10^. That work provided the first evidence of epigenetic dysregulation in primary antibody deficiencies (PADs) and has spearheaded interest in epigenetic aberrancies in the broader field of primary immunodeficiency research^11^. However, that study only interrogated a limited set of CpG sites and did not explore links with the functional impact of such epigenetic defects. DNA methylation has an established role in B cell differentiation and biology^12, 13^. It is likely that DNA methylation alterations are accompanied by other epigenetic changes and by transcriptional alterations in the B cell compartment of CVID patients.

Single-cell transcriptomics and epigenomics emerge as novel approaches for studying the diversity, plasticity and adaptability of immune cell subsets^14^. Here, we used single-cell atlas technologies to identify defects occurring specifically in scarce immune subsets that would otherwise be masked in conventional bulk-based approaches. By studying the aforementioned pair of CVID-discordant MZ twins and a cohort of patients and healthy donors, we generated and integrated single-cell multi-omics datasets (DNA methylation, chromatin accessibility and transcriptomics) to understand the heterogeneity and impact of epigenetic dysregulation in CVID.

Our data revealed prevalent and widespread alterations affecting DNA methylation and chromatin accessibility of memory B cells, and provided evidence that these changes impact the transcriptome of these cells following activation. Additionally, our single-cell transcriptomics census of whole blood cells revealed defective cell–cell communication, which may explain the establishment of aberrant epigenetic and transcriptional programs, as well as the impaired immune responses observed in these patients.

## RESULTS

### Genome-wide DNA methylation alterations in CVID occur during memory B cell differentiation

To define the DNA methylation profiles in CVID, we performed single-cell whole-genome bisulfite sequencing of B cells isolated from a pair of CVID-discordant MZ twins (Fig. 1a) without identified pathogenic mutations. Clinical information, vaccination response and B cell phenotype of these individuals are displayed in Supplementary Data 1. Specifically, we generated the whole DNA methylomes^15^ of around 200 single cells (Supplementary Data 2) corresponding to naïve, un-switched memory (US-mem) and switched memory (S-mem) B cells from the control and CVID twins (Fig. 1a and Supplementary Fig. 1 and Supplementary Fig. 2). We then imputed missing methylation values using DeepCpG^16^ per cell type, and generated merged data from the same subpopulations from each donor (pseudo-bulks) to outline their methylomes.

**Figure 1.**
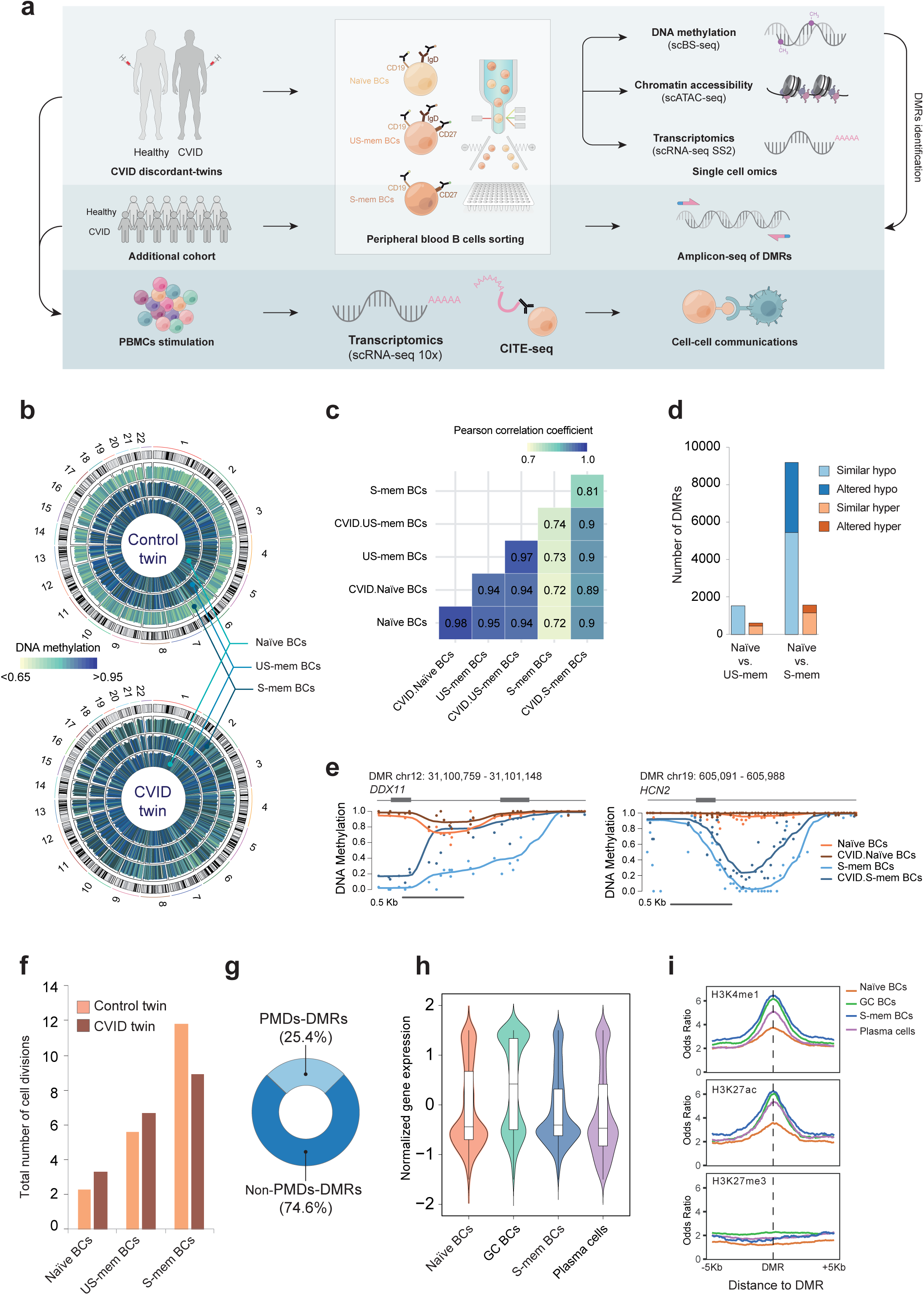
CVID methylation defects take place in the memory compartment at genomic regions involved in B cell function. (a) Scheme depicting sorter strategy for B cell isolation from the CVID-discordant twins and the cohort of CVID patients and healthy controls, as well as the multi-omics single-cell approaches used. (b) Circular representation of DNA methylation levels in naïve B cells, US-mem B cells, and S-mem B cells from control and CVID twins. Histogram tracks represent the average methylation levels over 5 Mb windows throughout the genome. Autosomal chromosomes from 1 to 22 are represented. (c) Heatmap showing Pearson correlation coefficients from the comparison between control and CVID within the various B cell compartments. (d) Bar plot representing the number of significant DMRs identified (q<0.05 and meth.diff>20%) in the transition from naïve to US-mem B cells or from naïve to S-mem B cells. (e) Selected examples of smoothed DNA methylation data in altered DMRs in the CVID twin. (f) Bar plot indicating the total number of cell divisions in naïve, US-mem and S-mem B cells in control and CVID twins. (g) Overlap of *CVID.no-demeth DMRs* with partially methylated domains (PMDs, light blue; non-PMDs, dark blue). (h) Violin plots representing normalized expression levels of the genes nearest to *CVID.no-demeth DMRs* in naïve, GC and S-mem B cells, as well as in PCs. Gene expression data (n=1 biological independent sample for each B cell population) were obtained from publicly available RNA-seq datasets (see Methods). Whiskers correspond with the minimum and maximum values of the data set (excluding any outliers). The box is drawn from Q1 to Q3 with a horizontal line to indicate the median. (i) Plots showing odds ratios in *CVID.no-demeth DMRs* ± 5 Kb flanking regions in naïve, GC and S-mem B cells, as well as PCs for different enhancer-associated histone modifications (H3K4me1, H3K27ac and H3K27me3). Active enhancer regions were defined for the concurrence of H3K4me1 and H3K27ac, repressed enhancers by H3K4me1 and H3K27me3, whereas primed enhancers by H3K4me1 exclusively. Histone modifications data were obtained from publicly available ChIP-seq datasets (see Methods).

In parallel, we performed whole genome sequencing (WGS) of the MZ twins to study a potential genetic origin of the CVID discordance observed in these individuals. WGS analysis revealed the existence of 1,400 somatic variants in the CVID twin (Supplementary Data 3), which is in the same range as the variants observed in twin cohorts^17, 18^. According to ClinVar^19^ there are only two differential SNPs and two differential indels between the twins in genes potentially related to a disease (<u>VCV000332641.2</u>, <u>VCV000670396.1</u>, <u>VCV000801645.1</u> and <u>VCV000402355.1</u>), although none of these genetic variants have previously been linked to CVID. We also analyzed the effect of the identified genetic variants using SNPEff^20^ and observed that only one of them induced a relevant effect on the gene *TAS2R31*. The link of this gene to CVID or to any other disease has not been described (Supplementary Data 3). Of note, we identified genetic variants shared by both CVID discordant twins affecting genes previously associated with CVID, such as *CR2*, *NFKB2*, *CD19* and *TNFRSF13B*. This observation reinforces the hypothesis that additional alterations, such as epigenetic changes, might be contributing factors to the differential phenotype of these siblings to the development of CVID.

In relation to the single-cell methylation analysis, in the control twin, DNA methylation levels of both US-mem and S-mem B cells decreased relative to naïve B cells (Fig. 1b, top, and Supplementary Fig. 3a), consistent with the progressive demethylation occurring during B cell differentiation^12, 13^. However, we found that S-mem B cells in CVID displayed impaired DNA demethylation throughout the entire genome (Fig. 1b, bottom, and Supplementary Fig. 3a). The methylomes of naïve and US-mem B cells from the CVID sibling were virtually identical to those of their control counterparts, whereas the methylomes of S-mem B cells displayed profound differences between the two twins, as ascertained by Pearson correlation and principal component analysis (PCA) (Fig. 1c and Supplementary Fig. 3b-3c).

To identify the functional loci potentially involved in transcriptional regulation, we defined differentially methylated regions (DMRs) using *Metilene*^21^ (Supplementary Data 4 and Supplementary Fig. 1). In the transition from naïve-to-US-mem B cells, we detected 1,537 DMRs that underwent hypomethylation in both control and CVID twins. However, of the 628 DMRs that became hypermethylated in control, 192 were not hypermethylated in CVID (Fig. 1d). The most dramatic alterations were found in the transition from naïve-to-S-mem B cells. Specifically, 3,745 of the 9,204 hypomethylated DMRs (40.7%) of the naïve-to-S-mem B cells comparison of the healthy twin, did not undergo demethylation in his CVID sibling. Of the 1,555 hypermethylated DMRs in the naïve-to-S-mem B cell differentiation in the healthy control, 404 (26%) were not hypermethylated in CVID (Fig. 1d and 1e). No DMRs were found in the comparison of naïve B cells from CVID and healthy control.

FACS analysis showed that the proportions of IgG+ and IgA+ B cells within the memory compartment of the twins are comparable (around 50% of each in both individuals), eliminating the possibility that altered methylation observed in the CVID twin was a consequence of a different memory B cell subtype abundance.

Our results indicate that CVID patients display methylation defects that do not preexist in naïve B cells but are established during memory B cell differentiation, and affect only specific genomic regions. Since most of the alterations detected in CVID affected DNA methylation loss and occurred in S-mem B cells, we focused on those regions and referred to them as *CVID.no-demeth DMRs* hereafter for brevity (Supplementary Data 5).

A high proportion of the DNA demethylation taking place during cell differentiation occurs in genomic regions named *partially methylated domains* (PMDs), which are characterized by late replication and passive demethylation^22, 23^. To test whether the impaired DNA demethylation observed in CVID S-mem B cells results from alterations in passive cell-division-associated demethylation, we analyzed their replicative history using the KREC assay^24^. Consistent with our previous findings^25^, S-mem B cells of the CVID twin displayed a lower cell division rate (Fig. 1f), which would be compatible with reduced levels of passive PMD-associated demethylation. However, the majority of the *CVID.no-demeth DMRs* overlapped with non-PMDs and with early replication genomic regions (Fig. 1g and Supplementary Fig. 3d), indicating a prevalence of alterations in active demethylation mechanisms during S-mem B cell differentiation in CVID. We found no significant differences in the expression of methylcytosine dioxygenase Ten-eleven translocation (TET) enzymes TET1, TET2 and TET3, involved in active demethylation, in the different B cell compartments (Supplementary Fig. 3e), suggesting a defect in the enzyme activity and/or its recruitment to genomic sites rather than an alteration in the levels of these proteins.

To evaluate the functional relevance of the *CVID.no-demeth DMRs*, we analyzed publicly available gene expression data from various B cell populations (accession numbers indicated in Methods). We observed that a large number of the nearest genes to these *CVID.no-demeth DMRs* were specifically upregulated in germinal center (GC) B cells compared with naïve B cells, S-mem B cells or plasma cells (PCs) from healthy individuals (Fig. 1h) and corresponded to functional categories related to B cell biology (Supplementary Fig. 3f). Additionally, the vast majority of the *CVID.no-demeth DMRs* were located at intronic and intergenic regions (Supplementary Fig. 3g), suggesting a role in modulating regulatory elements. In this regard, using publicly available ChIP-seq data of histone marks (accession numbers indicated in Methods), we observed that *CVID.no- demeth DMRs* were more highly enriched in active enhancer histone marks (H3K4me1 and H3K27ac) in GC B cells, S-mem B cells and PCs than in naïve B cells (Fig. 1i). All these results suggest that CVID aberrant hypermethylation in S-mem B cells occurs at genomic regions that control B cell function, comprising genes and enhancer regions that become activated during GC reaction.

### Heterogeneous aberrant DNA methylation patterns define key transcription factors associated with B cell activation in CVID

We assessed the heterogeneity of DNA methylation profiles in the S-mem B cell compartment from the CVID-discordant twins. PCA based on hypomethylated DMRs in the naïve-to-S-mem B cell transition revealed greater methylation heterogeneity in S-mem than in naïve B cells (Fig. 2a). Some of the cells within the CVID S-mem B cell compartment displayed similar DNA methylation profiles to those of control cells, whereas others clustered separately and showed aberrant hypermethylation (Fig. 2a and 2b). In-depth analysis of the data (see Methods) showed that the entire set of S-mem B cells from the CVID twin displayed methylation alterations, although the frequency of these alterations was highly variable (Fig. 2c).

**Figure 2.**
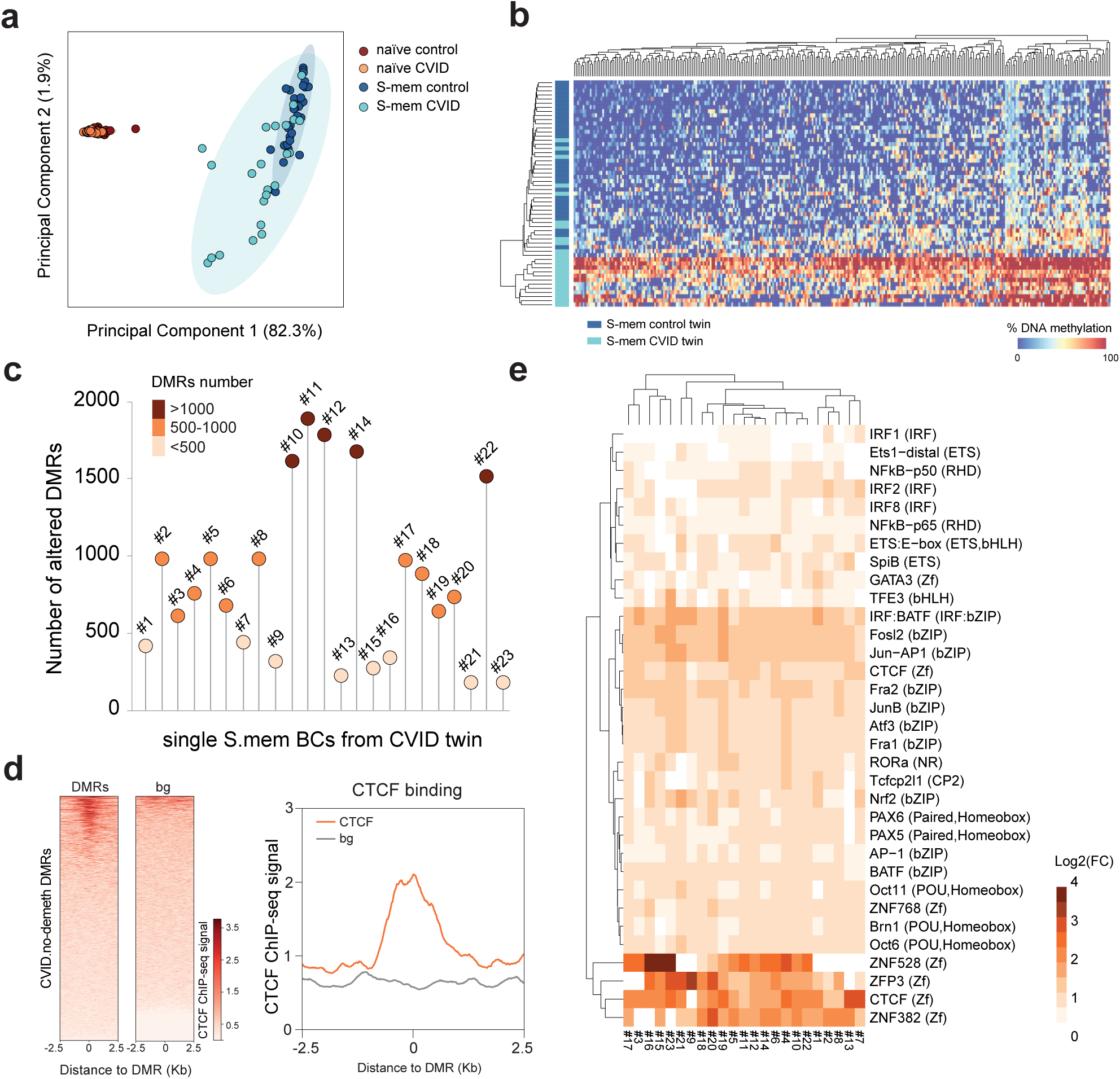
Heterogeneous DNA methylation alterations in CVID S-mem B cells. (a) Dot plot showing principal component analysis based on the DMRs at the single-cell level of naïve and S-mem B cells in the control and CVID twins. The elliptical shadows correspond to the confidence interval (90% CI) for each subpopulation. (b) Heatmap representing the DNA methylation percentage of the 300 most variable *CVID.no-demeth DMRs* at the single-cell level. Dark blue samples correspond to S-mem B cells from the control twin; light blue samples correspond to S-mem B cells from the CVID twin. (c) Plot showing the number of altered hypomethylated DMRs for each individual S-mem B cell for the entire set of cells analyzed at the single-cell level in the CVID twin. (d) Heatmap and histogram depicting CTCF binding signal in human primary B cells at *CVID.no-demeth DMRs* ± 2.5 Kb flanking regions. CTCF binding data was obtained from publicly available ChIP-seq dataset (see Methods). (e) Heatmap representing the log_2_(FC) enrichment for several TF binding motifs at the single-cell level over the altered hypomethylated DMRs in S-mem B cells from the CVID twin.

Transcription factor (TF) motif enrichment analysis of *CVID.no-demeth DMRs* revealed significant enrichment of several TFs involved in B cell differentiation and function, including members of the bZIP family (e.g., BATF, JUNB, Fosl2 and Fra2), NF-kB (p65 and p50), CTCF, IRFs and PU.1^21^ (Supplementary Fig. 4a). Using publicly available ChIP-seq data from human primary B cells and immortalized B cells (accession numbers indicated in Methods), we observed a greater degree of binding for several of these TFs, such as CTCF, BATF and JUNB in *CVID.no-demeth DMRs*, which implies that the regions are indeed bound to these TFs *in vivo* in B cells (Fig. 2d and Supplementary Fig. 4b). Motif enrichment analysis of *CVID.no-demeth DMRs* at the single-cell level showed similar motif enrichment for TFs, such as CTCF, NF-kB-p65, IRF8 and members of the bZIP family, in almost all of the individual cells, regardless of the number of altered DMRs they presented (Fig. 2e).

Our results indicate that, despite the heterogeneous demethylation defects affecting CVID S-mem B cells, the entire compartment displayed DNA methylation alterations in common TF binding sites that might compromise the proper binding of those TFs.

### Chromatin accessibility alterations occur in regulatory elements and are independent of methylation defects in CVID

In addition, we performed single-cell assay for transposase-accessible chromatin with sequencing (scATAC-seq) of the B cell compartment to determine chromatin accessibility profiles and explore their relationship with the previously identified DNA methylation alterations in CVID. Uniform Manifold Approximation and Projection (UMAP) visualization showed that the three B cell subsets analyzed (naïve, US-mem and S-mem B cells) were clustered on the basis of their chromatin accessibility status (Fig. 3a).

**Figure 3.**
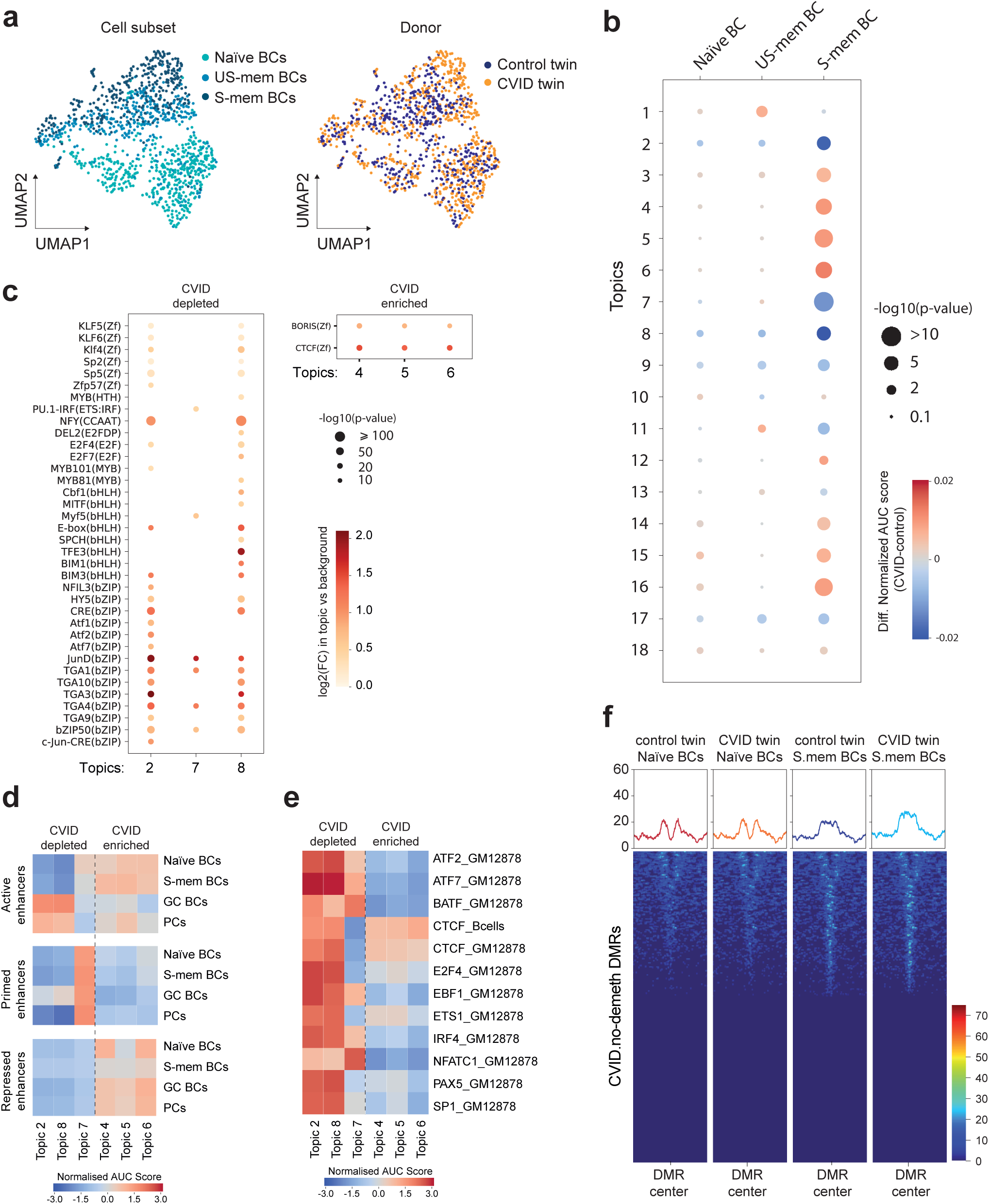
Chromatin accessibility defects in CVID. (a) UMAP visualization of unsupervised analysis of chromatin accessibility in naïve, US-mem and S-mem B cells from the control and CVID twin indicating the B cell subset and the origin of the cell. (b) Heatmap representing the difference in normalized AUC scores for the different regulatory topics defined by CisTopic between CVID and control twins in the B cell subsets analyzed. Red color corresponds to a stronger association of the topic with the CVID twin; blue corresponds to a stronger association of the topic with the control twin. Circle size indicates the significance of the association (Mann-Whitney test). (c) Dotplot representing the significantly enriched TF binding motifs in selected topics that are most closely associated with chromatin accessibility in control twin (CVID-depleted: topics 2, 7 and 8) or most associated with chromatin accessibility in the CVID twin (CVID-enriched: topics 4, 5 and 6). (d) Heatmap representing the association of the CVID-depleted and CVID-enriched topics with different enhancer stages during B cell activation (active, primed and repressed enhancers) Active enhancer regions were defined for the concurrence of H3K4me1 and H3K27ac, repressed enhancers by H3K4me1 and H3K27me3, whereas primed enhancers by H3K4me1 exclusively. (e) Heatmap showing the association of CVID-depleted and CVID-enriched topics with several TFs binding signal in human immortalized B cells and CTCF in human primary B cells. (f) Heatmaps and histograms representing chromatin accessibility signals in naïve and S-mem B cells at *CVID.no-demeth DMRs* ± 2.5 Kb flanking regions in control and CVID twins.

We then used *cisTopic*^26^ to identify chromatin accessibility modules (groups of co-occurring accessible chromatin regions referred to hereafter as topics) in scATAC-seq data for the three B cell subpopulations (Supplementary Fig. 1). A topic may be linked to a particular biological feature in the cell. As in the previous DNA methylation analysis, most CVID-associated differences in chromatin accessibility occurred in the S-mem B cell compartment (Fig. 3b). Among the 18 regulatory topics identified using *cisTopic*, we selected six of them with the greatest differences (above the 95^th^ and below the 5^th^ percentiles). Topics 2, 7 and 8 corresponded to less accessible regions in CVID S-mem B cells (CVID-depleted topics), whereas topics 4, 5, and 6 defined more accessible regions in CVID S-mem B cells (CVID-enriched topics) (Fig. 3b). TF binding motif analysis of these topics indicated the significant enrichment of several members of the bZIP, bHLH and ZF TF families in CVID-depleted topics, and enrichment of CTCF in CVID-enriched topics (Fig. 3c), coincident with several of the TF binding motifs enriched in the *CVID.no-demeth DMRs*.

Subsequently, we defined active (H3K4me1 and H3K27ac), primed (H3K4me1) and repressed (H3K4me1 and H3K27me3) enhancers occurring in naïve, S-mem and GC B cells as well as in PCs, and investigated how the identified topics were associated with those regulatory elements. We observed a stronger association of CVID-depleted topics with active enhancers in GC B cells and PCs (Fig. 3d). Our results indicate the existence of CVID chromatin accessibility defects in regulatory elements that undergo activation upon antigen encountering by naïve or memory B cells. Consistent with this observation, we detected greater motif enrichment of several TFs involved in enhancer regulation, such as BATF, IRF4 and CTCF^27–29^ in CVID-depleted topics (Fig. 3e).

In order to validate the findings obtained with cisTopic, we also performed an unsupervised clustering of the cells based on scATACseq data (Supplementary Fig. 4c). Of note, among the 9 identified clusters, cluster 3 (in red) was significantly enriched in CVID B cells compared with control B cells (Fisher’s test p=0.0001). This cluster also displayed the highest number (384) of significant differentially accessible regions (DARs, calculated with limma, p<0.05). Among those regions, 356 DARs were less accessible in the CVID sample (CVID-depleted), and 28 were more accessible (CVID-enriched). To further give those DARs meaning in terms of the previous features that we had inspected (Fig. 3d,e), we examined the overlap of TFs binding or enhancer regions (see Methods, using the same publicly available files as for Fig. 3d,e) with those DARs (Supplementary Fig. 4d,e). The analysis of this specific cluster 3 showed that CVID-depleted DARs were enriched in active enhancers in GC or post-GC states (memory B cells and plasma cells), which is in line with our previous findings using the cisTopic tool (Supplementary Fig. 4d). It also indicates that all the TFs peaks examined were more enriched in CVID-depleted DARs which is consistent with our previous observations for topics 2 and 8 in cisTopic analysis (Supplementary Fig. 4e). These results may suggest that chromatin accessibility dysregulation in the CVID twin is taking place in a specific subset of memory B cells.

We also noted an increase in chromatin accessibility in *CVID.no-demeth DMRs* in the naïve-to-S-mem B cell transition. However, those regions became accessible despite not being properly demethylated in the CVID twin (Fig. 3f). This analysis suggests that increased chromatin accessibility might be a prior process that is necessary but not sufficient for DNA demethylation and that, in the CVID twin, defects in the proper recruitment/activation of the demethylating machinery is impaired.

Overall, our results indicate that chromatin accessibility alterations in CVID S-mem B cells occur in regulatory elements that become active during the B cell response. Additionally, the impaired DNA demethylation observed in CVID proves not to be a consequence of changes in chromatin accessibility.

### CVID epigenetic alterations contribute to transcriptional perturbations in activated B cells

DNA methylation and chromatin accessibility defects in CVID S-mem B cells are likely to affect gene expression. We therefore performed plate-based Smart-seq2^30^ single-cell RNA-sequencing (RNA-seq) in naïve, US-mem and S-mem B cells from the aforementioned CVID-discordant twins. Unsupervised dimensionality reduction, clustering and differential expression analysis in each of the subsets revealed virtually identical transcriptional profiles in both twins (Fig. 4a, Supplementary Fig. 1 and Supplementary Data 6).

**Figure 4.**
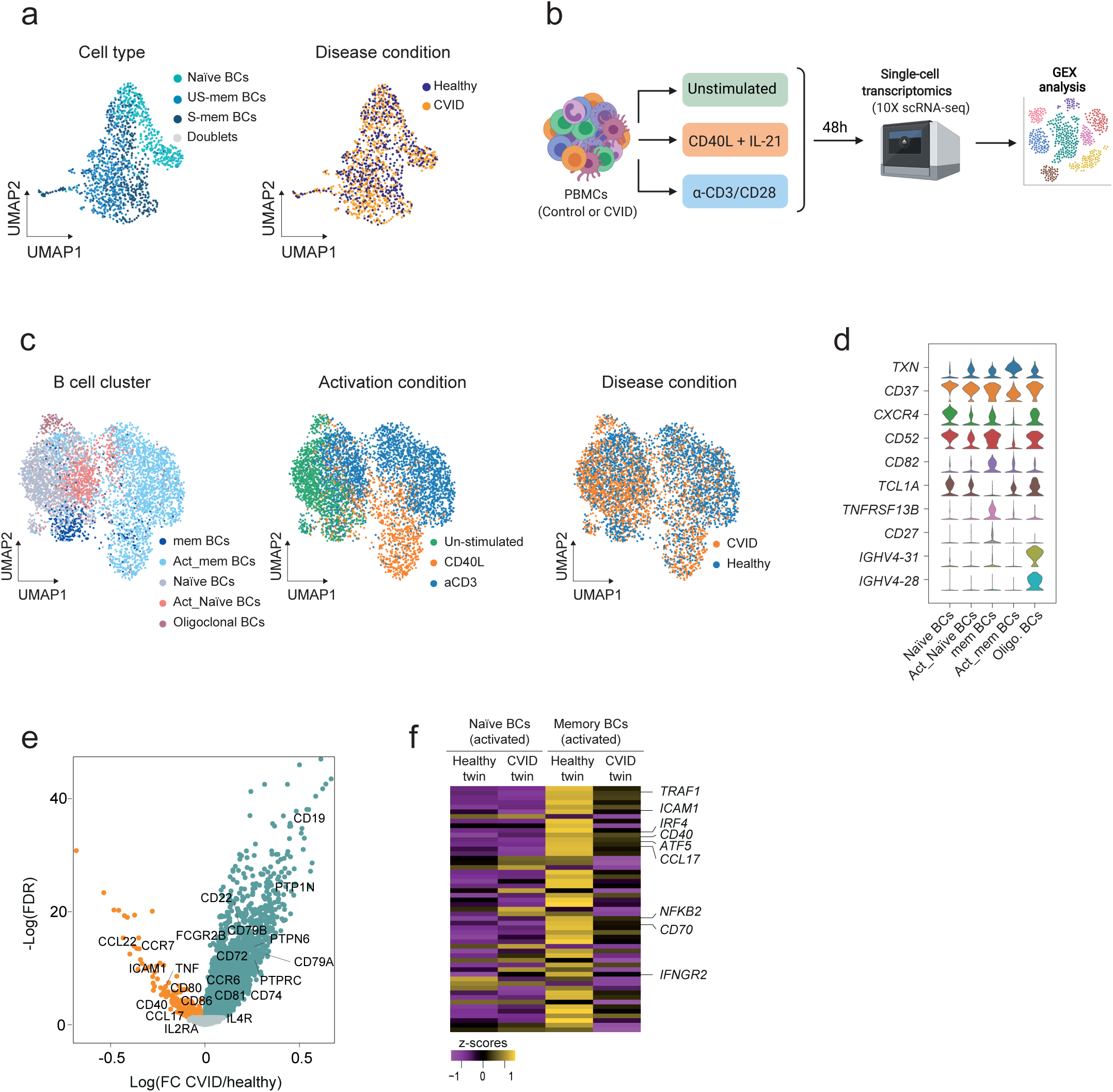
CVID transcriptional defects in activated B cells. (a) Unsupervised UMAP visualizations of SS2 single-cell transcriptome data based on highly variable genes and indicating B cell subset and cell origin. (b) Scheme depicting immune cell activation with CD40L + IL-21 (B cell activation) or α-CD3/CD28 (T cell activation), the establishment of cell–cell communications after cell stimulation and the use of 10x Chromium technology for single-cell transcriptomic analysis. (c) Unsupervised UMAP visualizations showing the predicted classification of B cells into steady state or activated naïve and memory B cells using logistic regression, the stimulation conditions and disease distributions. (d) Violin plots representing the expression of selected marker genes used to annotate the B cell subpopulations in the CVID and healthy twins (e) Volcano plot representing differentially expressed genes in activated memory B cells comparing control and CVID twins. Some relevant upregulated and downregulated genes are labeled. (f) Heatmap representing the expression levels of selected dysregulated genes in CVID activated memory B cells corresponding to the nearest gene to a CVID.no-demeth DMR. Activated naïve B cells and activated memory B cells from CVID discordant twins are represented.

The transcriptional signatures of naïve and memory B cells are very similar despite the profound epigenetic reprogramming of naïve B cells following antigen encounter^31^. We therefore speculated that epigenetic alterations in CVID might influence the transcriptome of activated B cells, instead of the transcriptome of steady-state B cells (naïve and memory B cells without any stimulus). This would also be consistent with our analysis showing that DNA methylation and chromatin accessibility alterations in CVID are both associated with activated B cell features (GC upregulated genes and GC active enhancers).

Therefore, we studied the transcriptome of activated B cells from the CVID-discordant twins. To this end, we conducted *in vitro* activation using peripheral blood mononuclear cells (PBMCs). Specifically, we stimulated PBMCs using a combination of CD40L and IL-21 (CD40L/21) (to directly activate B cells) or, alternatively, using α-CD3/CD28 (CD3/28) (to define the influence of the T cell compartment on B cells). We then used droplet-based techniques^32^ to create a transcriptomic census of different immune cells of the CVID-discordant twin pair and performed gene expression analysis (Fig. 4 and Supplementary Fig. 1).

Under these experimental conditions, cell deconvolution using logistic regression and cell markers expression analysis allowed us to identify naïve and memory B cell subsets (Fig. 4c and 4d and Supplementary Fig. 5a). Despite the high similarities between the control and the CVID individual in the expression of B cell activation marker genes (Supplementary Fig. 5b), differential expression analysis identified 870 upregulated and 11 downregulated genes (FDR < 0.05) in the activated naïve B cell cluster of the CVID twin (Supplementary Fig. 5c and Supplementary Data 7). However, in line with the previously identified epigenetic defects occurring specifically in the memory B cell compartment, we detected 6-fold more dysregulated genes in activated memory B cells (5,116 upregulated and 179 downregulated genes) when the CVID twin was compared with his healthy control sibling (Fig. 4e and Supplementary Data 7). Dysregulated genes in activated memory B cells included B cell-related upregulated genes such as *FCGR2B*, *CD72*, *PTPRC, CD79A, CD22, and CCR6*, as well as downregulated genes such as *CCL22*, *CD40*, *ICAM1*, *CCR7, CCL17, CD80* and *CD86* (Fig. 4e).

Separate differential gene expression analysis of the two activation conditions (CD40L + IL-21 or anti-CD3/CD28), identified that relevant genes such as *CCL22*, *IL4R*, *IRF4*, *ATF5* and *IFNGR2* were specifically dysregulated upon CD40L + IL-21 activation, whereas *CD74*, *CD79A*, *CD79B*, *PTPN6*, *CD72*, *CD22* or *PTPN1* were dysregulated specifically upon anti-CD3/CD28 stimulation. Interestingly, other genes such as *CD19*, *LGALS9*, *TGFB1*, *PTPRCAP* and *LY6E* were similarly dysregulated regardless of the activation condition (Supplementary Data 7). However, we noticed that separate analysis causes a drastic decrease in the statistical power of the analysis with a reduction of 50 % in the number of detected differentially expressed genes. In this regard, a large group of the previously identified DEGs such as *PTPRC*, *NFKB2*, *TNF*, *CD80*, *CD86*, *CD40*, *CCR6* and *CD70*, among others, were not detected as differentially expressed now (Supplementary Data 7). The dysregulation of relevant genes for B cell activation such as *CD79A*, *CD79B*, *CD22*, *CD72* and *PTPN6*, all of them genes involved in the modulation of the B cell receptor pathway, specifically upon anti-CD3/CD28 treatment (indirect B cell activation via T cell stimulation) and not upon CD40L + IL-21 treatment (direct B cell stimulation), would suggest the existence of extrinsic B cell defects that might contribute to the impaired B cell responses observed in CVID patients.

As shown above, most of the DNA methylation alterations in the CVID twin occur in enhancers active during the B cell response. However, we identified a group of altered genes in CVID activated memory B cells that were also in the proximity of aberrantly hypermethylated regions (Fig. 4f). 907 (18%) of the upregulated genes and 53 (30%) of the downregulated genes in S-mem BCs of the CVID twin correspond to genes near *CVID.no-demeth DMRs*. Interestingly, some of these hypermethylated and downregulated genes were specifically upregulated in memory B cells upon cell activation in comparison with naïve B cells. For instance, selected members of the CD40 signalling pathway (eg. *CD40*, *TRAF1*, and *NFKB2*), and other genes related to B cell response (eg. *CD70*, *ICAM1* (CD54), *CCL17*) were specifically hypermethylated and downregulated in activated memory B cells in the CVID twin (Fig. 4f).

To integrate the multiple single-cell -omic data generated in this study (i.e. scBS-seq, scATAC-seq and scRNA-seq data), we firstly analyzed the expression of relevant TFs. We then related the concurrence of the dysregulated TF with the presence of TF motifs in the regions with DNA methylation and chromatin accessibility alterations. We identified that the expression levels of members of the IRF, ETS and bZIP, TF families such as IRF1, SPIB, JUN and FOS were significantly altered in the activated memory B cells of the CVID twin (Supplementary Data 7). Interestingly, several members of those TF families were also significantly enriched in the genomic regions displaying epigenetic alterations (Supplementary Fig. 5d), both in DNA methylation (previously shown in Fig. 2e) and in chromatin accessibility (previously shown in Fig. 3e).

Our findings demonstrate that the transcriptional program of activated B cells is compromised in the CVID twin and that CVID-associated epigenetic alterations might contribute to transcriptional perturbations that occur during B cell response.

### CVID is characterized by immune cell communication defects mediating B cell responses

Our single-cell transcriptomics census allowed us to generate the transcriptional profiles of immune compartments additional to B cells from the CVID-discordant twin pair. Clustering and cell type-specific marker gene expression allowed us to identify several populations of immune cells: B cells (including naïve and memory B cells), CD4+ and CD8+ T cells (naïve and memory), CD8+ MAI T cells, Treg cells, γδ-T cells, two subsets of NK cells, myeloid cells and their corresponding activated subsets (Fig. 5a-5b and Supplementary Fig. 6a). Differential gene expression analysis between the two CVID-discordant siblings within each compartment revealed transcriptional dysregulation in immune subsets other than B cells (Supplementary Data 8). In this regard, these immune compartments displayed dysregulation of pathways related to the IFN signature, as previously described in CVID^33, 34^ (data not shown).

**Figure 5.**
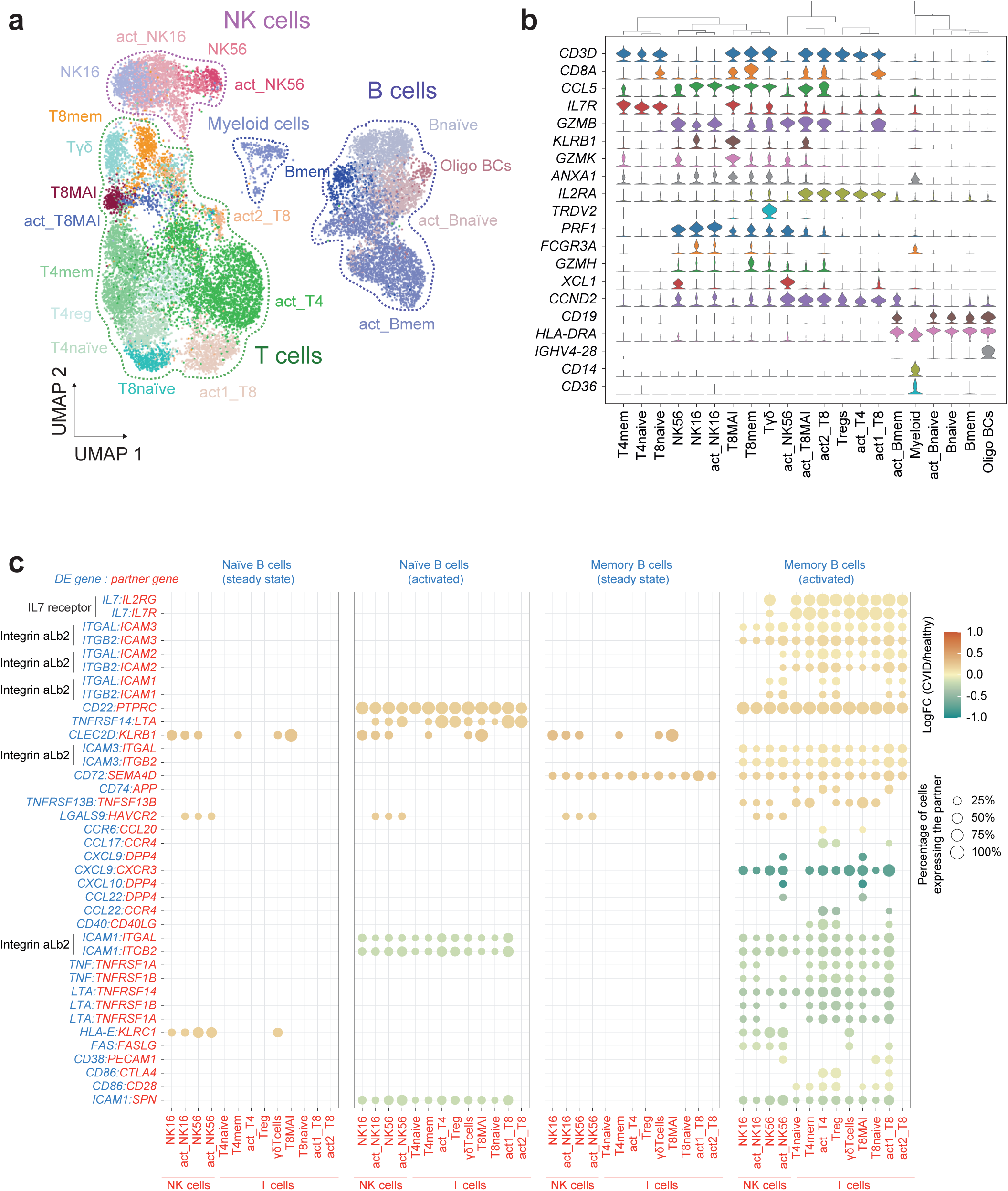
Cell–cell communication alterations in the CVID twin. (a) UMAP visualization showing different immune cell populations identified from Louvain clustering and cell-specific marker gene expression. The B cell compartment is outlined with a blue dotted line and includes naïve BCs (Bnaïve), memory BCs (Bmem) activated naïve BCs (act_Bnaïve), activated memory BCs (act_Bmem) and oligoclonal BCs (oligo BCs). The T cell compartment is outlined with a green dotted line and includes CD8+ naïve T cells (T8naïve), CD8+ memory T cells (T8mem), CD8+ activated T cells (act1_T8 and act2_T8), CD8+ MAI T cells (T8MAI), CD8+ activated MAI T cells (act_T8MAI), CD4+ naïve T cells (T4naïve), CD4+ activated T cells (act_T4), CD4+ regulatory T cells (Treg) and γδ-T cells (Tγδ). The NK cell compartment is outlined with a purple dotted line and includes NK CD16+ cells (NK16), activated NK16 cells (act_NK16), NK CD56+ cells (NK56) and activated NK56 cells (act_NK56). Myeloid cells are also represented (b) Violin plots representing the expression of selected marker genes in the cell populations identified. (c) Overview of selected dysregulated L/R interactions between B cells and the other immune cell compartments in the CVID twin in naïve and memory B cells in steady state and following activation. The scale indicates the log_2_(FC) gene expression of the defined B cell subsets in the CVID *vs.* control comparison. Only differentially expressed genes with an FDR < 0.05 were considered in the analysis. The percentage of other immune cells expressing the partner molecule is indicated by the circle size. Molecules of the L/R pairs expressed in B cells are shown in blue; molecules of the L/R pairs expressed in other non-B immune cells are shown in red. Assays were carried out at the mRNA level, but were extrapolated to protein interactions.

To systematically analyze the effect of cell–cell communication on B cell responses in the healthy and the CVID twins, we used CellPhoneDB^35^ (Supplementary Fig. 1), our recent database of ligands, receptors and their interaction. This approach, available on GitHub (https://github.com/Teichlab/cellphonedb), allowed us to generate a potential cell–cell communication network upon immune cell challenge in the CVID-discordant twins.

CellPhoneDB analysis revealed defects in several ligand-receptor (L/R) pairs relevant to B cell regulation (Fig. 5c and Supplementary Data 8). Remarkably, activated memory B cells displayed the highest number of dysregulated L/R pairs (Fig. 5c and Supplementary Fig. 6b). For instance, several genes that code for negative regulators of the BCR function, such as CD22, CD45 (*PTPRC*), Galectin-9 (*LGALS9*) and CD72, were significantly upregulated in CVID B cells, which might explain the defective B cell response in CVID twins. The interacting partners of these upregulated genes in B cells were expressed in several immune cells that are present both in the NK and T cell compartments (Fig. 5c). Furthermore, we observed that the negative regulators CD22 and CD45 were also upregulated in CVID B cells, which might constitute a repressive autocrine/paracrine mechanism affecting B cell activation in the CVID twin (Supplementary Fig. 6b). Conversely, we also observed that several genes, like *CD86* and the chemokine genes *CXCL10*, *CCL22* and *CCL17*, were significantly downregulated in CVID B cells. These cytokines presented their receptor partners in both NK and T cell subsets (Fig. 5c). We also found expression changes in other cell compartments that interact with B cells (Supplementary Fig. 6c and Supplementary Data 8). For instance, in the CD4+ T cell compartment, we observed significant downregulation of CD40L, the ligand of CD40, which is expressed in B cells and is required for B cell activation. This was also observed at the protein level by FACS (Supplementary Fig. 6d), although due to the inherent limitations of these particular rare samples, this is a single experiment that should not be taken as definitive proof.

All these results indicate that, in addition to B cell-intrinsic alterations, defects in other immune cell compartments might compromise the correct B cell response in the context of this primary immunodeficiency.

### DNA methylation, transcriptomic and cell-cell communication defects in a CVID cohort

We performed DNA methylation and transcriptomic analysis in a cohort consisting of 10 CVID patients and an equivalent number of healthy controls and compared the results obtained in our twin-study. The CVID patients -without any known monogenic defect-represented the three CVID subtypes (Ia, Ib and II) according to the Freiburg classification ^36^. Clinical information and B cell phenotype of these individuals are provided in Supplementary Data 1.

We first performed amplicon sequencing to interrogate 1058 CpG sites corresponding to 162 selected *CVID.no-demeth DMRs* (Supplementary Fig.1 and Supplementary Data 9). We observed that the main methylation alterations observed in the CVID discordant twin are also present in the validation cohort (Fig. 6a). Selected DMRs (Fig. 6b), confirmed the occurrence of hypermethylation in S-mem B cells. Methylation levels in S-mem B cells were significantly higher for all 3 CVID subtypes represented in our cohort (Fig. 6c). Interestingly, the effects were most accentuated for those CVID patients of the Ib subtype (the subtype to which the CVID twin belongs). Furthermore, in agreement with our previous works^10, 25^, we observed aberrant hypermethylation in genes such as *BCL2L1, TCF3, BCL6, BCL10*, and *AICDA* among others (Supplementary Fig. 7a). As in the case of the twins (Fig. 1f), CVID patients showed lower proliferation rates in both naïve and memory B cells after CD40L + IL-21 stimulation (Supplementary Fig. 7b). The existence of these proliferative alterations in the naïve B cell compartment raises the possibility of additional primed defects established in previous steps of B cell development.

**Figure 6.**
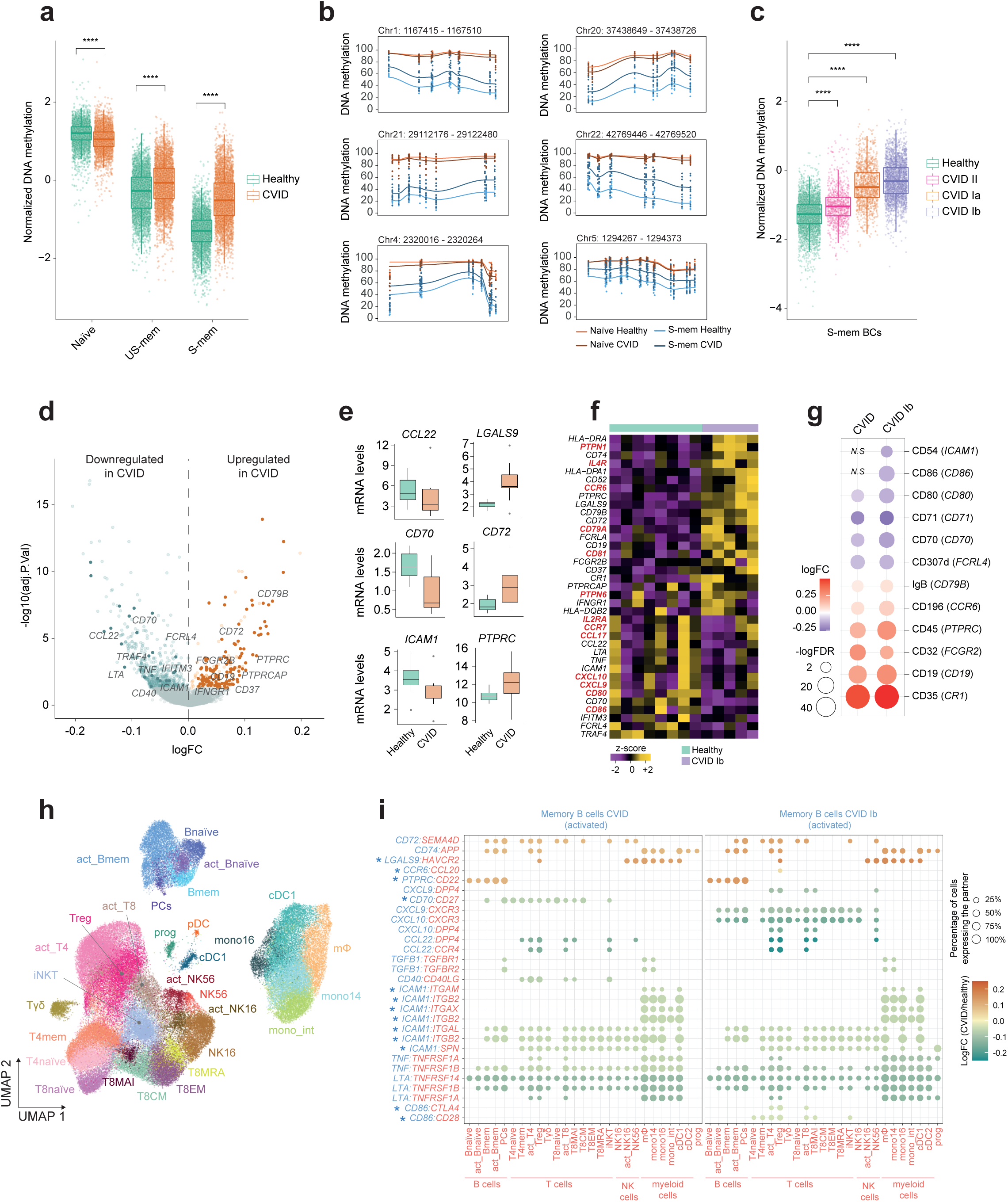
DNA methylation, transcriptomic and cell-cell communication defects in a CVID cohort. (a) Box-plot depicting DNA methylation levels of selected DMRs in different B cell subsets from CVID patients (n=11) and healthy individuals (n=10). Whiskers correspond with the minimum and maximum values of the data set (excluding any outliers). The box is drawn from Q1 to Q3 with a horizontal line to indicate the median. Two-sided Wilcoxon test with no multiple test correction, where **** represents p-value <= 0.0001. (b) Plot showing smoothed DNA methylation data in altered DMRs . (c) Box-plots depicting DNA methylation levels of the inspected CpG sites in S-mem B cells from CVID patients (n=11) and healthy controls (n=10). Two-sided Wilcoxon test with no multiple test correction, where **** represents p-value < 0.0001. (d) Volcano plot representing DEGs in activated memory B cells comparing healthy donors (n=8) and CVID patients (n=10). Downregulated (green) and upregulated (orange) genes are plotted. Dysregulated genes in the CVID-discordant twin were highlighted with a darker color. R elevant genes are labeled. (e) Box-plots representing selected dysregulated genes in the CVID cohort (n=10) compared to healthy donors (n=8). (f) Heatmap representing DEGs from the comparison of healthy controls and CVID Ib patients. Genes also dysregulated in the discordant twins comparison are labeled in red. (g) Dot plot representing selected differentially expressed proteins (gene name in brackets). S cale indicates the log_2_(FC) protein expression of the activated memory B cell subset in the CVID or CVID Ib *vs* control comparison, -logFDR is indicated by the circle size. *N.S* means not significant. (h) UMAP visualization showing different immune cell populations identified. (i) Overview of selected dysregulated L/R interactions. Scale indicates the log_2_(FC) gene expression of the memory activated B cell subset in the CVID or CVID Ib *vs* control comparison.. The percentage of other immune cells expressing the partner molecule is indicated by the circle size. Molecules of the L/R pairs expressed in activated memory B cells are shown in blue; molecules of the L/R pairs expressed in the immune cell partner are shown in red. Validated gene expression at the protein level is indicated with an asterisk.

We also obtained the single-cell transcriptomes of steady-state and activated B cells in the cohort of CVID patients and healthy controls. We obtained the transcriptome of ∼100K PBMCs stimulated with (i) either CD40L and IL-21 (CD40L/21), or (ii) α-CD3/CD28 (CD3/28) (Supplementary Fig. 1 and Supplementary Fig. 7c). We observed shared transcriptomic signatures between the CVID-discordant twin pair and the new CVID cohort. Within the B cell compartment, activated memory B cells displayed the highest number of dysregulated genes, as previously observed for the CVID twin (Fig. 6d-6e and Supplementary Data 10). Several of these differentially expressed genes in both the CVID cohort and the CVID twin, include relevant genes for B cell immune response such as *CCL22*, *CD70*, *ICAM1*, *LGALS9*, *CD72* or *PTPRC*. Inspection of the CVID Ib subcohort led us to identify additional dysregulated genes shared with the CVID twin. Examples of these genes include *CCR6, PTPN1, CD81 and CD79A* (upregulated) *and CCR7, CD80, CD86 and CCL17* (downregulated) (Fig. 6f and Supplementary Data 11). Some of these dysregulated genes were validated at the protein level using CITE-seq analysis (Fig. 6g and Supplementary Tables 12, 13 and 14). For example, we detected protein downregulation of CD54 (*ICAM1*), CD86, CD80, CD70, CD71 and CD307d (*FCRL4*), together with protein upregulation of IgB (*CD79B*), CD196 (*CCR6*), CD45 (*PTPRC*), CD32 (*FCGR2*), CD19 and CD35 (*CR1*) in the activated memory B cells of the CVID patients compared with the healthy donors. In addition, the analysis of a larger number of cells in the CVID cohort allowed us to annotate additional immune cell populations such as progenitors, DCs and monocytes among others, besides the ones identified in the twin analysis (Fig. 6h and Supplementary Fig. 7d). We also observed shared dysregulated ligand-receptor pairs in the cohort analysis in comparison to the twins dataset not only in the B cell compartment (eg. *CD72:SEMA4D, CD74:APP, CCL22:CCR4, LGALS9:HAVCR2* and *CD40:CD40LG* among others) but also in compartments beyond B cells such as activated CD4+ T cells (Fig. 6i, left panel, Supplementary Fig. 7e and Supplementary Fig. 8a-b) As previously observed for DEGs, we detected additional dysregulated L/R pairs when focusing in CVID Ib subtype patients (eg. *CCR6:CCL20, CXCL9:CXCR3, CXCL10:DPP4* and *CD86:CTL4* among others) (Fig. 6i, right panel and Supplementary Data 15).

## DISCUSSION

In this study, the use of multi-omics single-cell atlas technologies allowed us to identify functional alterations occurring in specific scarce immune cell populations that would have gone undetected if conventional bulk-based approaches had been used. We show that CVID B cells display widespread heterogeneous DNA methylation and chromatin accessibility changes that are restricted to the memory compartment. Similarly to other epigenomic studies for congenital syndromes^37, 38^, the DNA methylation patterns identified in this study might lead to the identification of a diagnostic “episignature” for CVID. Epigenetic signatures in the S-mem B cell compartment can be used as a historical record of epigenetic processes occurring during the GC reaction. In this regard, we have demonstrated that epigenetic alterations in CVID memory B cells affect genes and regulatory elements that undergo activation in GC B cells and subsequent PC differentiation.

DNA demethylation changes in CVID S-mem B cells are mainly associated with altered active demethylation mechanisms, as shown by the predominance of alterations outside PMDs. We did not detect any significant changes in the expression levels of ten-eleven translocation (TET) demethylating enzymes. However, given that several TFs interact with TETs^39–42^, it is likely that dysregulation of certain TFs might alter the recruitment of demethylating activities to specific genomic loci.

Regardless of the heterogeneous degree of CVID-associated methylation defects observed in the S-mem B cell compartment, we detected a common enrichment in the binding motif of TFs associated with B cell function in those altered regions. For instance, Fra1 is involved in the regulation of follicular B cell differentiation into PCs^43^. In addition, IRF8 is crucial to the development of GC B cells and to the maintenance of the B cell program^44, 45^, and CTCF is required to sustain the GC transcriptional program to avoid premature PC differentiation^46^. Interestingly, several studies have shown how DNA methylation may affect the binding of some TFs, including CTCF, AP-1 and BATF^47–49^. Therefore, the observed methylation defects may compromise the functionality of the entire memory B cell compartment in CVID.

CVID-associated epigenetic alterations are not reflected in the transcriptomes of the few steady-state S-mem B cells that are present in peripheral blood and which are virtually identical to those in the healthy controls. Instead, B cells display transcriptional differences upon activation. Since several donors included in this study are healthy individuals, any invasive approach including a lymph node biopsy would not be ethically justifiable for research purposes. Given these constraints, we used *in vitro*-activated B cells in order to partially emulate the GC reaction. The GC reaction is a tightly regulated process in which the balance between survival and apoptotic signals is critical to promote antigen affinity while simultaneously avoiding self-antigen recognition. In this context, we identified significant downregulation of several chemokine genes in CVID B cells such as *CCL22* and *CCL17*, which have been described as fine-tuning antibody affinity maturation and positive selection in GC promoting B cell interactions with follicular helper T cells^50^. Other relevant downregulated genes in the B cells of CVID patients are those encoding CD70, which has been involved in B cell activation ^51, 52^, and ICAM1 (CD54), whose interaction with the integrin LFA-1 lowers the threshold for B cell activation by promoting B cell adhesion and immunological synapse formation^53^. Additionally, our data indicate that genes encoding Gal-9, CD45 and CD22 are upregulated in the B cell compartment of the CVID twin. Gal-9 hampers BCR activation through its interaction with the N-glycan repertoire of CD45 molecules, which ultimately inhibits BCR signaling via CD22^54^. Another study showed that Gal-9 facilitates the interaction of IgM-BCR with the inhibitory molecules CD45 and CD22^55^. Additionally, we observed significant upregulation of the *CD72* gene in CVID B cells, which is another molecule that induces cell cycle arrest and apoptosis in mature B cells and that is involved in the inhibition of BCR downstream signaling pathways^56, 57^. The aberrant upregulation of these inhibitory molecules might interfere with the appropriate activation of the BCR, and might be one of the causes of the deficient immune responses and the defective B cell counts that characterize CVID patients.

It is of particular note that we also detected transcriptional defects beyond the B cell compartment. In this regard, we found several dysregulated genes in the activated CD4+ T cells of the CVID patients. For instance, we found downregulation of genes such as *SEMA4D* (CD100), which is an interacting partner of CD72, as well as upregulation of the B cell inhibitory molecule *PTPRC* (CD45). Additionally, we found a significant downregulation of *CD40LG* (CD40L) in this T cell compartment as previously described by others ^58, 59^, although the dysregulation of this molecule is still a contentious topic in the field^60^. All these alterations taking place in other immune subsets distinct to B cells, together with the specific dysregulation in the B cell compartment upon indirect B cell activation through T cell stimulation, would constitute B cell-extrinsic defects that also might contribute to the failure of B cell activation in CVID patients.

In addition to the cohort of CVID patients and healthy controls used, one of the strong points of this study is the use of monozygotic twins, which are genetically identical. It is likely that the exposure to different environments at prenatal or post-delivery time, or even their differential early infection history, may have induced epigenetic alterations that ultimately lead to a difference in gene expression and twin discordance in CVID. The reduction of both memory B cell and PC counts in CVID patients, as well as the severe hypogammaglobulinemia in one of the twins could be a consequence of the defects in the proper activation of naïve B cells, alterations during memory and PC generation, or even both. However, our results indicate that naïve B cell populations are almost identical in both twins in terms of their DNA methylation, chromatin accessibility and transcription profile, indicating that CVID epigenetic and gene expression defects are established later on during naïve B cell activation and memory B cell generation at secondary lymphoid organs, where the establishment of appropriate cell–cell communication is crucial for mounting efficient immune responses.

## METHODS

### Patients and ethics statement

Human blood samples used in this study were obtained from a pair of monozygotic twins discordant for CVID (the same pair participating in a previous study^10^ and an additional cohort of 10 CVID patients and 10 healthy donors (Supplementary Data 1). CVID patients were diagnosed according to European Society for Immunodeficiencies (ESID) criteria ^61^. They were collected at the University Hospital Dr Negrín, Gran Canaria, Spain and at the Hospital La Paz, Madrid, Spain. All donors received oral and written information about the possibility that their blood would be used for research purposes, and any questions that arose were then answered. Before giving their first blood sample the donors signed a consent form approved by the Ethics Committee at their corresponding hospital (Hospital La Paz PI-2833), which adhered to the principles set out in the WMA Declaration of Helsinki. The protocol used to isolate B cells from these donors was approved by the Ethics Committee of the Bellvitge University Hospital (CEIC) on 9 March 2017 (PR053/17).

### Sample collection and immune cell activation

PBMCs were obtained from peripheral blood by Ficoll gradient using LymphoprepTM (Stemcell Technologies. Cat. No. # 07801). For the isolation of CD19+ CD27^neg^ IgD+ naïve, CD19+ CD27+ IgD+ unswitched memory (US-mem) and CD19+ CD27+ IgD^neg^ switched memory (S-mem) B cells, PBMCs were stained with anti-CD19-FITC (BD Biosciences , clone: 4G7. Cat. No. 15856028), anti-CD27-APC (Miltenyi Biotec, clone: M-T271. Cat. No. 130-113-630) and anti-IgD-PE (Southern Biotech, Birmingham, AL, USA, clone: NA, Ref: 2032-09) in MACS buffer (PBS + 2% FBS + 2 mM EDTA).

For immune cell activation, PBMCs were resuspended in complete medium (RPMI Medium 1640 + GlutaMAX^TM^ (Gibco, Life Technologies. Cat. No. 61870036) containing 20% fetal bovine serum (Gibco, Life Technologies. Cat.No. 10270-106), 100 units/mL penicillin, and 100 μg/mL streptomycin). PBMCs were then activated with a B cell stimulus consisting of a combination of 0.1 µg/mL of CD40L (MEGACD40L® Protein, ENZO. Cat. No. ALX-522-110-C010) and 50 ng/mL of IL-21 (Tebu-bio. Cat. No. 200-21), with a T cell stimulus consisting of α-CD3/CD28 Dynabeads 20 μL of beads/million PBMCs (Invitrogen. Cat. No. 11131D) for 48 hours. As a control, some PBMCs were also cultured with no stimulus. After stimulation, PBMCs were harvested and a fraction of the cells were stained with anti-CD19-FITC for B cell sorting in Beckman Coulter MoFlo Astrios EQ Cell Sorter instrument. Finally, PBMCs were mixed with sorted CD19+ B cells in a 2:1 ratio in order to obtain PBMCs samples enriched in B cells. This B cell enrichment step was performed in order to avoid T cells and other immune cell types mask the B cell compartment due to the drastically reduced levels of B cells observed in CVID patients.

For the analysis of CD40L expression in CD4+ T cells, PBMCs were isolated from the CVID discordant twins and cultured at a concentration of 200,000 cells/200 uL of complete medium in a 96 well plate. Cells were stimulated with α-CD3/CD28 Dynabeads 20 μL or 4 μL of beads/million of PBMCs for 48 hours. As a control, some PBMCs were also cultured with no stimulus. After stimulation, PBMCs were harvested and stained with anti-CD4-APC (BD Biosciences, clone: RPA-T4. Cat. No. 555349) and anti-CD40L-PE (BD Biosciences, clone: TRAP1. Cat. No. 555700) in MACS buffer. Cells were acquired using Gallios Flow Cytometer (Beckman Coulter, CA, USA) and analyzed using the Flowjo software.

### Whole genome sequencing analysis

Whole genome sequencing (WGS) of genomic DNA obtained from the pair of monozygotic twins discordant for CVID. The quality of the isolated genomic DNA was checked on 1% agarose gel. Besides, DNA concentration was measured using Qubit® DNA Assay Kit (Cat. No. 10146592) in Qubit® 2.0 Fluorometer (Life Technologies, CA, USA). The genomic DNA was randomly fragmented by sonication to the size of 350 bp, then DNA fragments were end polished, A-tailed, and ligated with the full-length adapters of Illumina sequencing, and followed by further PCR amplification. The PCR products as the final construction of the libraries were purified with the AMPure XP system. Then libraries were checked for size distribution using Agilent 2100 Bioanalyzer (Agilent Technologies, CA, USA), and quantified by real-time PCR (to meet the criteria of 3 nM).

The 150×2 paired-end sequencing was conducted on a Novaseq sequencer (Illumina, CA, USA) producing 612,180,910 raw paired reads on average.

Quality check (QC) and read alignment were performed using miARma-seq pipeline^62^. This software uses FastQC (version 0.11.5)^63^ to detect adapter contamination and low quality reads. Reads with a quality lower than 30 were stringently eliminated to reduce false positives from the sequencing output. After QC, the sequence data were subjected to BWA (version 0.7.13)^64^ for alignment to the human reference genome (GRCh38p12). The alignment data (BAM files) were processed by GATK (version 4.1.7.0)^65^ according to their best practices workflows. In that sense markduplicates, base recalibration, covariate analysis and variant filtration (for SNPs: Variant Confidence/Quality by Depth (QD)< 2.0, Phred-scaled p-value using Fisher’s exact test to detect strand bias (FS) > 60.0, RMS Mapping Quality (RMQ)< 40.0, Symmetric Odds Ratio of 2×2 contingency table to detect strand bias (SOR) > 4.0, Z-score From Wilcoxon rank sum test of Alt vs. Ref read mapping qualities (MQRankSum) < -12.5 and Z-score from Wilcoxon rank sum test of Alt vs. Ref read position bia (ReadPosRankSum) < -8.0. For indels: QD < 2.0, FS > 200.0, SOR > 10.0) was performed to optimize the final depth and base-call quality for variant detection.

The set of common variants between both individuals was obtained using the intersectBed tool from the bedtools suite^66^. Furthermore, putative somatic mutations were detected with the MuTect2 software by GATK (version 4.1.7.0) by comparing BAM files from the twin and the co-twin. A base recalibrator process was performed providing known sites of variation such as the allele frequency by the Genome Aggregation Database (gnomAD) ^67^, high confidence SNPs and indels from the 1000 Genomes project^68^ and the NCBI database of genetic variation (dbSNP)^69^, to label sequencing errors and distinguish poor base quality from variants. These fine-tuned parameters allowed to optimize the final depth and base-call quality for somatic mutation Identification. Then, the FilterMutectCalls tool from the GATK toolbox was applied to use allele fractions to distinguish somatic variants from sequencing errors.

The relationship of identified variants with phenotypes was performed with the intersectBed utility using the data provided by the ClinVar database^19^. The impact of the sequence modification by these variants at the functional level was annotated using SNPEff^20^.

### KREC assay

The replication history of naïve, US-mem and S-mem B cells from CVID MZ discordant twins, was inferred from isolated DNAs by the κ−deleting recombination excision circle (KREC) assay, as previously described^24^. The replication history is estimated by the ratio between genomic coding joints and signal joints on KRECs of the Igκ intron RSS-K deleting rearrangement, which are quantified by Real-Time PCR, applying the formula: ΔCT sample= Ct (signal joint) - Ct (coding joint).

### Single-cell library preparations

For single-cell bisulfite sequencing (scBS-seq) assays, naïve, US-mem and S-mem B cells were collected by sorting in 12 μl of lysis buffer (10 mM Tris-Cl pH 7.4, 0.6% SDS, 0.5 μl proteinase K) using a Beckman Coulter MoFlo Astrios EQ Cell Sorter instrument in single-cell 1 drop mode. ToPro-3 and Hoechst 33342 staining were used to select for live cells with low DNA content (i.e., in G0/G1). Negative controls were sorted using BD Accudrop Beads on lysis buffer, and were prepared and processed concomitantly with all single-cell samples. Libraries for scBS-seq were generated as previously described^15^ but with the following modifications. Bisulfite conversion was performed using an Imprint DNA modification kit (Sigma. Cat. No. MOD50-1KT). Purification and desulphonation of converted DNA was performed with magnetic beads (Zymo) on a Bravo Workstation (Agilent), eluting into the master mix for the first strand synthesis. Purified scBS-seq libraries were sequenced in pools of 12-14 single-cell libraries were prepared for 100bp paired-end sequencing on a HiSeq2500 in rapid-run mode (2 lanes/run).

The plate-based single-cell ATAC-seq assay was performed as previously described^70^. Briefly, naïve, US-mem and S-mem B cells were collected by FACS in MACS buffer, and cells were centrifuged at 500*g* at 4 °C for 5 min. Cell pellets were resuspended in 50 μl tagmentation mix (33 mM Tris-acetate, pH 7.8, 66 mM potassium acetate, 10 mM magnesium acetate, 16% dimethylformamide, 0.01% digitonin and 5 μl of Tn5 from the Nextera kit from Illumina, Cat. No. FC-121-1030). The tagmentation reaction was done on a thermomixer at 800 rpm, 37 °C for 30 min. The reaction was then stopped by adding equal volume (50 μl) of tagmentation stop buffer (10 mM Tris-HCl, pH 8.0, 20 mM EDTA, pH 8.0) and left on ice for 10 min. A volume of 200 μl MACS buffer with 0.5% BSA was added and the nuclei suspension was transferred to a FACS tube. DAPI was added at a final concentration of 1 μg/μl to stain the nuclei. DAPI-positive single nuclei were sorted into 96-well full-skirted Eppendorf plates chilled to 4°C, prepared with 2 µl 2X lysis buffer (100 mM Tris.HCl, pH 8.0, 100 mM NaCl, 40 µg/ml Proteinase K (Ambion, AM2546, 20 mg/ml stock) and 0.4% SDS) and 2 µl of 10 µM S5xx/N7xx Nextera Index Primer Mix (5 µM each). Plates were immediately sealed, spun down at 300*g* at 4°C for 1 min and stored at -80°C.

For single-cell RNA-seq using the plate-based Smart-seq2 (SS2) protocol, naïve, S-mem and S-mem B single cells were collected in 96-well full-skirted Eppendorf plates chilled to 4°C, prepared with lysis buffer consisting in 10 μl of TCL buffer (Qiagen. Cat. No. 1070498) supplemented with 1% β-mercaptoethanol. Plates were sealed, vortexed, spun down at 300 *g* at 4°C for 1 min, immediately placed on dry ice and transferred for storage at −80°C. The Smart-seq2 protocol was performed basically as previously described^71^. Libraries were sequenced, aiming for an average depth of 1 million reads per cell, on an Illumina HiSeq 2000 with version 4 chemistry (paired-end, 75-bp reads).

For the droplet scRNA-seq method, stimulated and unstimulated PBMCs enriched with CD19+ B cells were counted using a Neubauer hemocytometer, and 5,000 cells per sample were loaded in the 10x-Genomics Chromium. The 10x-Genomics 5’ libraries were prepared following the manufacturer’s instructions. Libraries were sequenced, aiming at a minimum coverage of 50,000 raw reads per cell, on an Illumina HiSeq 4000 (paired-end; read 1: 26 cycles; i7 index: 8 cycles, i5 index: 0 cycles; read 2: 98 cycles). For samples of the expanded cohort of CVID patients and healthy controls, we processed PBMC samples for scRNAseq with paired measurements of 192 surface proteins (CITE-seq, using TotalSeq reagents from Biolegend). The list of the antibodies included in the TotalSeq custom panel are detailed in Supplementary Data 12. Briefly, 1 million cells pooled from several individuals were stained with TotalSeq-C custom human panel together with Human TruStain FcX (Biolegend. Cat. No. 422301) on ice for 30 minutes. Then cells were washed three washes with cold PBS + 4% FBS. After completing the last washing steps, cells were resuspended in PBS + 0.04% BSA and filtered with a 40 um Bel-art Flowmi strainer into a new 1.5 ml lo-bind eppendorf tube. Cells were counted and concentration adjusted to load 50,000 cells on the 10X Genomics Chromium Station. The 10x-Genomics 5’ libraries together with CITE-seq libraries were prepared following the manufacturer’s instructions.

### Single-cell whole-genome bisulfite sequencing

For DNA methylation data, single-cell bisulfite sequencing (scBS-seq) data was processed as previously described^72^. Reads were trimmed with TrimGalore! (https://www.bioinformatics.babraham.ac.uk/projects/trim_galore/), Cutadapt^73^ and FastQC (https://www.bioinformatics.babraham.ac.uk/projects/fastqc/), using default settings for DNA methylation data and additionally removing the first 6 bp. Subsequently, Bismark^74^ (v0.16.3) was used to map the bisulfite data to the human reference genome (GRCh38), in single-end non-directional mode, followed by de-duplication and DNA methylation calling using default settings. We removed cells with a library size of fewer than 3 M reads, resulting in 196 cells with DNA methylation information. Specifically, there were 55 S-mem (23 CVID), 69 US-mem (31 CVID) and 72 naïve cells (37 CVID) (Supplementary Data 2).

### Imputation of DNA methylation signals

To mitigate the incomplete coverage of scBS-seq profiles, [healthy twin → naïve BCs= 27.0% ± 3.5%, US-mem BCs= 23.8% ± 5.9%, S-mem BCs= 21.8% ± 4.0% // CVID twin → naïve BCs= 23.0% ± 2.9%, US-mem BCs= 25.7% ± 8.5%, S-mem BCs= 24.2% ± 7.9%] (see Supplementary Data 2 for coverage percentages per single cell), we applied DeepCpG^16^ to impute unobserved methylation states of individual CpG sites. DNA methylation profiles of the three cell types were imputed separately; the two donors were jointly imputed to mitigate potential imputation artifacts between the samples. The cell type-specific models were developed using CpG and genomic information according to DeepCpG’s setup of a joint model (see DeepCpG^16^ for details and default values). The accuracy of the models per cell, as measured using AUC, ranged between 0.78 and 0.93, see Supplementary Data 2 for imputation accuracy per sample. Of note, the DNA methylation patterns after imputation recapitulates better the methylation profiles obtained from WGBS than those before imputation (when compared to data from the Blueprint consortium; EGAD00001002361 and EGAD00001002354), indicating that imputation is an appropriate approach to mitigate the technical noise associated with scBS.

### DMR identification

After imputation we merged the data per cell type - donor combination to identify differentially methylated regions (DMRs), using Metilene^21^ (v0.2.5) in *de novo* mode. We selected only DMRs with values of q < 0.05 and a methylation difference of > 20% for subsequent analysis. To identify altered DMRs in the CVID twin, we analyzed the previously selected DRMs using the limma package^75^, comparing the methylation values of those DMRs in single cells (FDR < 0.05 and methylation difference > 10%).

To define altered DMRs at the single-cell level in CVID, we first calculated the methylation distributions of each DMR in healthy control S-mem B cells. We then considered as altered DMRs those DMRs in CVID S-mem B cells whose mean methylation was more than 1.5*IQR above the third quartile or below the first quartile of the control DMR distributions.

### Amplicon sequencing for DNA methylation analysis

B cell subsets (naïve, US-mem and S-mem B cells) from 10 healthy donors and 11 CVID patients were sorted as described in the *Sample collection* section. Genomic DNA was then isolated from each population using QIAamp DNA micro kit (QIAGEN. Cat. No: 56304) and was bisulfite converted using MethylCode™ Bisulfite Conversion Kit (Thermofisher. Cat. No. MECOV50) following manufacturer instructions. From 1-10 ng of bisulfite converted DNA was used for amplicon generation using a custom panel, as well as for library preparation. The list of analyzed CpG sites (1058 CpG sites corresponding to 162 selected *CVID.no-demeth DMRs*) is available in Supplementary Data 9. Libraries were sequenced using IonTorrent technology (Thermofisher) according to manufacturer instructions. For methylation measure, only those CpG sites with a coverage > 50 (methylated + un-methylated reads) were considered.

### Analysis of the different -omics datasets in relation with several biological features

Transcription factor motifs were enriched for each set of identified CVID.hypo.DMRs in pseudo-bulk or in each single-cell using HOMER software v4.10.3. Specifically, we used the findMotifsGenome.pl algorithm (with parameters -size given -cpg) to search for significant enrichment compared with background, adjusted to ensure an equal number of sequences with equal lengths and with equal CpG and GC content.

DMR annotation for genetic context location was performed using the annotatePeaks.pl algorithm in the HOMER software v4.10.3.

Replication timing data in the GM12878 and GM12801 lymphoblastoid cell lines were obtained from the UW Repli-seq track of the UCSC Genome Browser. Replication timing values were binned in deciles to perform the overlap with the *CVID.no-demeth DMRs*.

GREAT software^76^ was used to enrich downstream pathways and gene ontologies. We used the single nearest gene option for the association between genomic regions with genes.

To define the genomic coordinates corresponding to active, primed or repressed enhancers, we used the following command via bedtools (version v2.28.0): “bedtools intersect file_A.bed file_B.bed”, in order to identify the regions with specific histone marks: For active enhancers file_A.bed corresponded to H3K4me1 peak file of the cell type of interest, and file_B.bed to H3K27ac. For the repressed enhancers file_A.bed corresponded to H3K4me1 peak file of the cell type of interest, and file_B.bed to H3K27me3. For defining primed enhancers, H3K4me1 bed file of the cell type of interest was used. All of the above was done separately for naïve, S-mem, GC B cells, as well as for plasma cells.

Inference of TF activities from expression values were calculated using DoRothEA^77^. The B_viperRegulon.rdata dataset was used for the analysis.

### Alignment, quantification and quality control of scRNA-seq data

Smart-seq2 (SS2) sequencing data were aligned with STAR^78^ (version 2.5.1b) using the GRCh38 (Ensembl release 84) STAR index and annotation. Gene-specific read counts were calculated using HTSeq-count (version 0.10.0). Cells with fewer than 200 detected genes and more than 20% mitochondrial gene expression content were removed. Genes expressed in fewer than 3 cells were removed.

Droplet-based RNA sequencing data for B cell receptor and T cell receptor (what we later on refer to as “BCR data” and “TCR data”) were aligned with Cell Ranger VDJ SingleCell Software Suite (version 3.0.2, 10x Genomics Inc.) using the GRCh38 official Cell Ranger V(D)J reference, version 2.0.0. Downstream analysis was performed using scirpy (https://icbi-lab.github.io/scirpy/).

Droplet-based RNA sequencing data was aligned and quantified using the Cell Ranger SingleCell Software Suite (version 3.0.2, 10x Genomics Inc.) using the GRCh38 human reference genome (official Cell Ranger reference, version 1.2.0). Cells with fewer 200 detected genes or more than 20% mitochondrial gene expression content were removed. Genes expressed in fewer than 3 cells were removed. For the SS2 and droplet-based datasets, the Scanpy (version 1.4.4) Python package^79^ was used to load the cell-gene count matrix and perform downstream analysis. To remove cell-cycle-associated variation, cell cycle-associated genes were removed from SS2 and droplet-based datasets. To do so, we normalized (*scanpy.pp.normalize_per_cell* method, scaling factor 10000), log-transformed (*scanpy.pp.log1p*) and subsetted the highly variable gene (*scanpy.pp.filter_gene_dispersion*) count matrices of both datasets. Next, datasets were transposed to operate in gene space with cells as features (gene-centered analysis). Data-feature scaling (*scanpy.pp.scale*), PCA analysis (*scanpy.pp.pca*, from variable genes), neighborhood graph building (*scanpy.pp.neighbors*) and Louvain graph-based clustering (*scanpy.tl.louvain*, clustering resolution manually tuned) were performed. Previously known cell cycle genes (*CDK1*, *MKI67*, *CCNB2*, *PCNA*) were used, and all genes that cluster together with any of these were removed.

### Deconvolution of donors in pooled CITE-seq samples

Pooled CITE-seq samples containing cells from multiple individuals were demultiplexed using souporcell ^80^. Briefly, the algorithm identifies genotypic differences between single cells by variant calling aligned reads using STAR ^78^ and generating a VCF (Variant Call Format) file using Freebayes (Marth 2012). Souporcell was run with the following command for all samples (argument $1 corresponds to sample ID, and N is the number of multiplexed individuals in a sample):

/software/singularity-v3.5.1/bin/singularity exec ./souporcell.sif ./souporcell_pipeline.py -i

./cellranger302_count_$1_GRCh38-1_2_0/possorted_genome_bam.bam -b

./cellranger302_count_$1_GRCh38-1_2_0/filtered_feature_bc_matrix/barcodes.tsv -f ./refdata-cellranger-GRCh38-1.2.0/fasta/genome.fa -t 8 -o souporcell_result_$1 -k N --skip_remap True -- common_variants ./filtered_2p_1kgenomes_GRCh38.vcf, where the last VCF file with common variants was downloaded as per instructions in https://github.com/wheaton5/souporcell with the following command: wget --load-cookies /tmp/cookies.txt “ https://docs.google.com/uc?export=download&confirm=$(wget --quiet --save-cookies/tmp/cookies.txt --keep-session-cookies --no-check-certificate

“https://docs.google.com/uc?export=download&id=13aebUpEKrtjliyT9rYzRijtkNJVUk5F_’ -O- | sed - rn ’s/.*confirm=([0-9A-Za-z_]+).*/\1\n/p’)&id=13aebUpEKrtjliyT9rYzRijtkNJVUk5F_” -O common_variants_grch38.vcf && rm -rf /tmp/cookies.txt.

Furthermore, to find out the exact donor identity of each donor’s barcode cluster in souporcell results, the cells in parallel have undergone genotyping using Illumina Infinium Global Screening Array. In order to unambiguously identify every individual in the pooled samples, each donor’s variants were separated from the pooled VCF and each single-donor VCF was matched to the genotype data using PLINK^81^. This software matches each souporcell sample with the genotype data giving a concordance ratio (based on the similarity of the variants) that allows us to distinctly identify each sample with each donor ID.

### Doublet removal from scRNA-seq and CITE-seq data

To exclude doublets from single-cell RNA and CITE sequencing data, a previously described two-step approach^82^ was used. Firstly, we applied the Scrublet^83^ algorithm for each sample to calculate the scrublet-predicted doublet score per cell. Then, to ensure that small clusters with high doublet density were not grouped with large numbers of singlets, we overclustered the manifold (running a basic scanpy pipeline up to clustering, additionally clustering each cluster separately), computed per-cluster Scrublet scores as the median of the observed values, and computed normal distribution parameters, centered on the median of the scores, with a MAD-derived standard deviation (the score distribution is zero-truncated, so we used only above-median values to compute the MAD). We then multiplied the MAD value by 1.4826 to obtain a literature-derived normal distribution standard deviation estimate and FDR-corrected the p-values using the Benjamini-Hochberg method^62^. Secondly, once the dataset had been processed and clustered with the Louvain algorithm as described above, step 1 was repeated for each of the clusters identified, thus capturing doublets within each cluster that were not detected with the initial doublet calling. Cells called as doublets in this last step were flagged as ‘Doublets’. Furthermore, we identified physical doublets of T and B cells using BCR data as cells in T cell clusters that had BCR data and, conversely, using TCR data as cells in B cell clusters that had TCR data. These cells were flagged as ‘B:T_doublets_by_TCR’ or ‘T:B_doublets_by_BCR’. In the zoomed in reanalysis of B cell populations of the twins we also labelled a cluster of cells expressing CD3E as ‘T_contaminants’. All the three types of detected doublet populations were removed from subsequent downstream analysis. In addition, for pooled samples, any cells that were called as inter-individual doublets by souporcell were also removed from subsequent downstream analysis.

### Denoising protein counts in CITE-seq data

To remove ambient signal from protein counts of the CITE-seq data, we used SoupX[https://academic.oup.com/gigascience/article/9/12/giaa151/6049831] in the following way: first, we got the expression profile of the ambient signal from empty droplets (estimateSoup with soupRange=c(4,100) ), then we automatically calculated contamination fraction (autoEstCont with soupQuantile= 0.1 and tfidfMin = 0.05) and removed background contamination from the count matrix (adjustCounts). The corrected protein counts were then merged with the gene expression part of CITE-seq data and further treated as raw counts and analysed. Protein features in the CITE-seq data were considered equal to gene features in subsequent downstream analysis.

### Clustering and annotation of scRNA-seq data and CITE-seq data

Downstream analysis included data normalization (scanpy.pp.normalize_per_cell method, scaling factor 10000), log-transformation (scanpy.pp.log1p), variable gene detection (scanpy.pp.filter_gene_dispersion), data-feature scaling (scanpy.pp.scale), PCA analysis (scanpy.pp.pca, from variable genes), batch-balanced neighborhood graph building (scanpy.pp.bbknn) by experimental sample batch key ‘sample’ and Louvain graph-based clustering (scanpy.tl.louvain, clustering resolution manually tuned). Cluster cell identity was assigned by manual annotation using known marker genes (Fig. 5b).

While annotating B cell populations, we perform more resolved clustering of the coarse “B cells” cluster (in the main manifold with all cells) to tease apart subtler B cell states (represented in Fig. 4c left panel) via (1) examining genes defining those states (as shown in Fig. 4d) and (2) examining logistic regression predictions from (a) unstimulated B cell scRNA-seq of twins (used in the first part of study and visualised in Fig. 4a) and (b) dataset from King et. al., Science Immunology, 2021 to better tell apart naïve and memory phenotype.

### Differential gene expression analysis in scRNA-seq

To calculate differentially expressed genes in each comparison cell state between CVID and control twin we adopted the limma^75^ approach (limma version 3.46.0, edgeR version 3.32.1).

### Cell–cell communication analysis

Cell-cell interactions were inferred using CellPhoneDB (www.CellPhoneDB.org), as previously described^35, 84^. We modified the statistical method to calculate significant L/R interactions enriched in disease. To do this, instead of random shuffling of cells to select specific interactions, we used genes differentially expressed between the twins in the different identified immune cell subsets (FDR < 0.05) and only included an L/R pair in the analysis when both molecules were present in at least 10% of the interacting cell clusters and at least one of the molecules was DE between CVID and healthy controls. Additionally, since CVID is considered a B cell-centered disease, we focused our analysis on the potential interactions between B cells and the other immune cell compartments.

### Alignment, quantification and downstream analysis of single-cell ATAC-seq data

Plate-based single-cell ATAC sequencing data were aligned and quantified as previously described^70^ and further analyzed with the standard scanpy pipeline described above (excluding cell cycle genes removal and doublet detection). Further downstream analysis was performed using *cisTopic*^24^ approach following the basic tutorial (https://rawcdn.githack.com/aertslab/cisTopic/f628c6f60918511ba0fa4a85366ebf52db5940f7/vignettes/CompleteAnalysis.html).

Transcription factor motif-enrichment analysis was performed for each identified topic using HOMER software v4.10.3. We used the findMotifsGenome.pl algorithm (with parameter -size given) to search for significant enrichment relative to the background.

Differentially accessible region (DAR) calculation was performed in a scRNA-seq like manner using limma ^75^ approach as described above. We used .bed files corresponding to significant DARs and .bed files corresponding to TF and enhancer Chip-seq data (same publicly available data as used for Fig. 3e,d) to first calculate overlaps between them and then estimate the enrichment of target (TF or enhancer) peaks in DARs as the ratio of total overlap length to length of target peaks.

### Data availability

All the data generated in this study as well as interactive visualizations of single-cell transcriptomic datasets from this study can be accessed via https://single-cell-atlas-cvid.cellgeni.sanger.ac.uk/. WGS results are not publicly available due to ethical regulations, but are available from the corresponding authors upon request.

RNA-seq and ChIP-seq publicly-available datasets for naïve-, GC, S-mem B cells, as well as for PCs from healthy donors were obtained from the Blueprint Consortium (http://dcc.blueprint-epigenome.eu/#/datasets), with the following accession numbers: Naïve BCs RNAseq (EGAD00001002315), GC BCs RNAseq (EGAD00001002452), S-mem BCs RNAseq (EGAD00001002476), Plasma cells RNAseq (EGAD00001002323), Naïve BCs ChIPseq (EGAD00001002466), GC BCs ChIPseq (EGAD00001002442), S-mem BCs ChIPseq (EGAD00001002430), Plasma cells ChIPseq (EGAD00001002281).

We used additional ChIP-seq publicly-available datasets for several transcription factors with the following accession numbers: CTCF from human B cells (GSM1003474), as well as JUNB (GSE96455), BATF (GSM803538), CTCF (ENCSR184YZV) and PAX8 (GSE127505) from the GM12878 human lymphoblastoid cell line.

### Code availability

All the code used in the analyses can be found at https://github.com/ventolab/CVID.

## Acknowledgements

We thank Antonio Gómez-García for graphical design support, Sarah Teichmann for her useful feedback, Hamish King for helping with single-cell germinal center dataset availability, Xi Chen for performing scATAC-seq analysis, Kirsty Ambridge and Elena Prigmore for their involvement in single-cell RNA library generation, Martin Prete for creating online visualizations for our cell atlas and Esther Castaño and Beatriz Barroso from CCiTUB Cytometry Unit for their support with single-cell sorting and Dr. Carla Gianelli and Dr. Rebeca Rodríguez Pena for the patient follow-up in the CVID cohort.

## Author contributions

J.R.-U., R.V.-T. and E.B. conceived the study; J.-R.U., L.P.-M. and L.C. designed and performed cell sorter, cell activation and KREC experiments; J.R.-U., L.C. and H.H. performed 10x Chromium experiments; J.R-U., S.J.C. and L.M. performed sample and library preparation; J.R-U., A.A., M.J.B, C.C.-F. L.G.-A., L.-F. H., E.A.L. F.K., K.P., S.V.D. and V.K. performed computational analysis; J.R.-U., A.A., J.M., K.W., R.V.-T. and E.B. analyzed and interpreted the data with contributions from L.P.-M, F.C.-M., V.R.-C., E.L.-G., C.R.-G., O.S. and G.K.; L.P.-M., M.M-S., E.L.-G. and C.R-G. contributed with clinical material and clinical interpretation of the results; J.R-U. and E.B. wrote the manuscript with contributions from A.A and R.V-T.; R.V.-T and E.B. co-directed the study. All authors read and accepted the manuscript.

