## Supplementary Materials for "Single-Cell Atlas of Common Variable Immunodeficiency reveals germinal center-associated epigenetic dysregulation in B cell responses"

### Supp. Figure 1

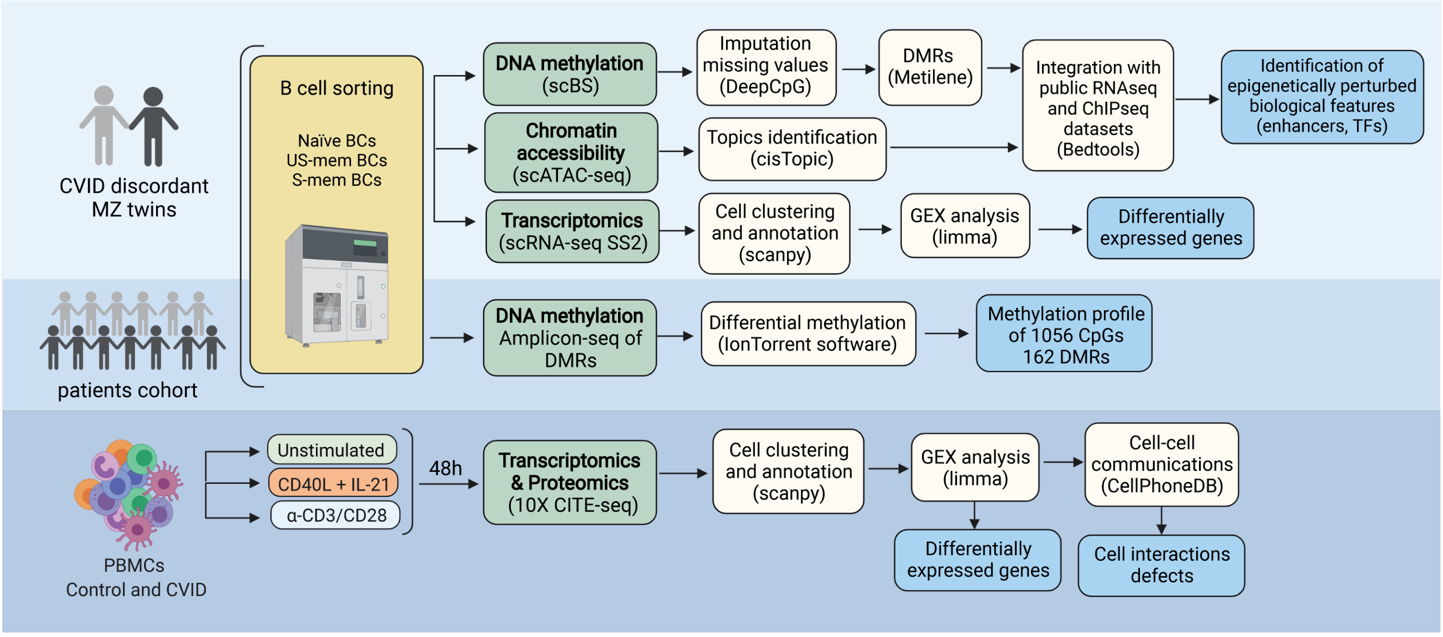

Supp. Figure 2

a

b

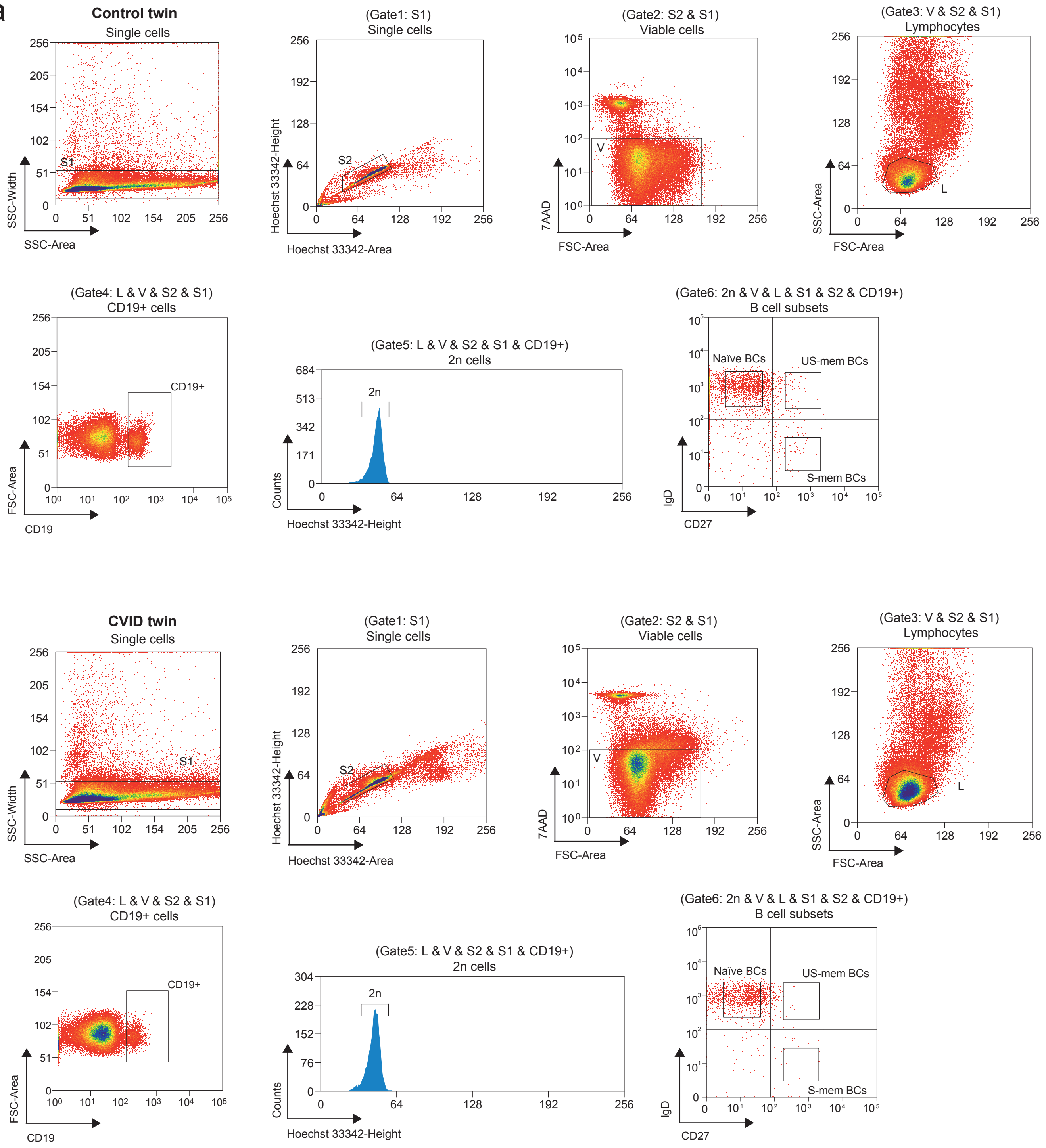


c

| Control twin | Abs (cells/uL) | % of PBL | % of CD19+ |
|---|---|---|---|
| Naïve | 145 | 3.20% | 75.10% |
| US-mem | 16 | 0.36% | 8.27% |
| S-mem | 16 | 0.37% | 8.57% |

| CVID twin | Abs (cells/uL) | % of PBL | % of CD19+ |
|---|---|---|---|
| Naïve | 191 | 4.10% | 78.10% |
| US-mem | 38 | 0.82% | 15.70% |
| S-mem | 1.4 | 0.03% | 0.59% |

### Supp. Figure 3

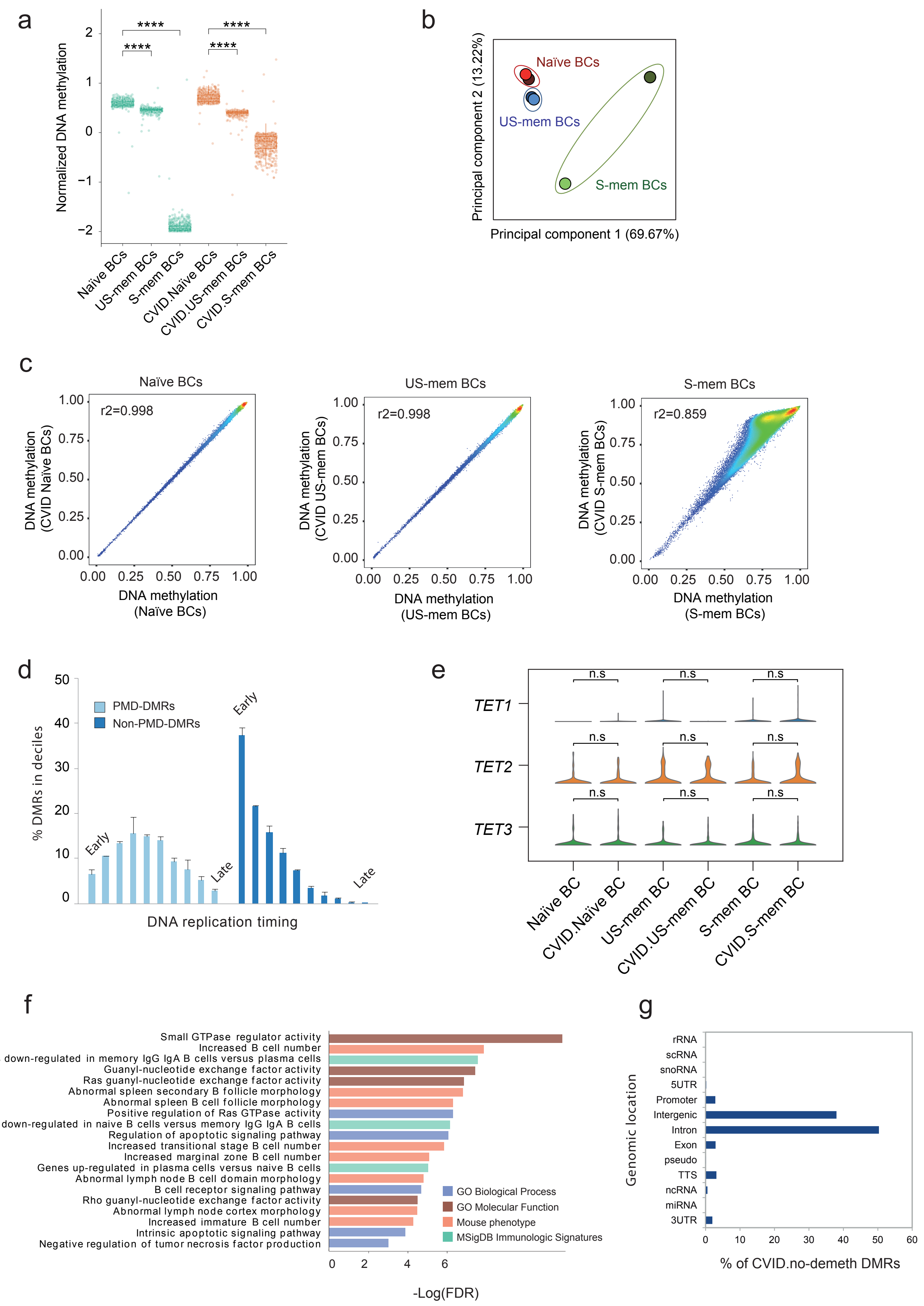

### Supp. Figure 4

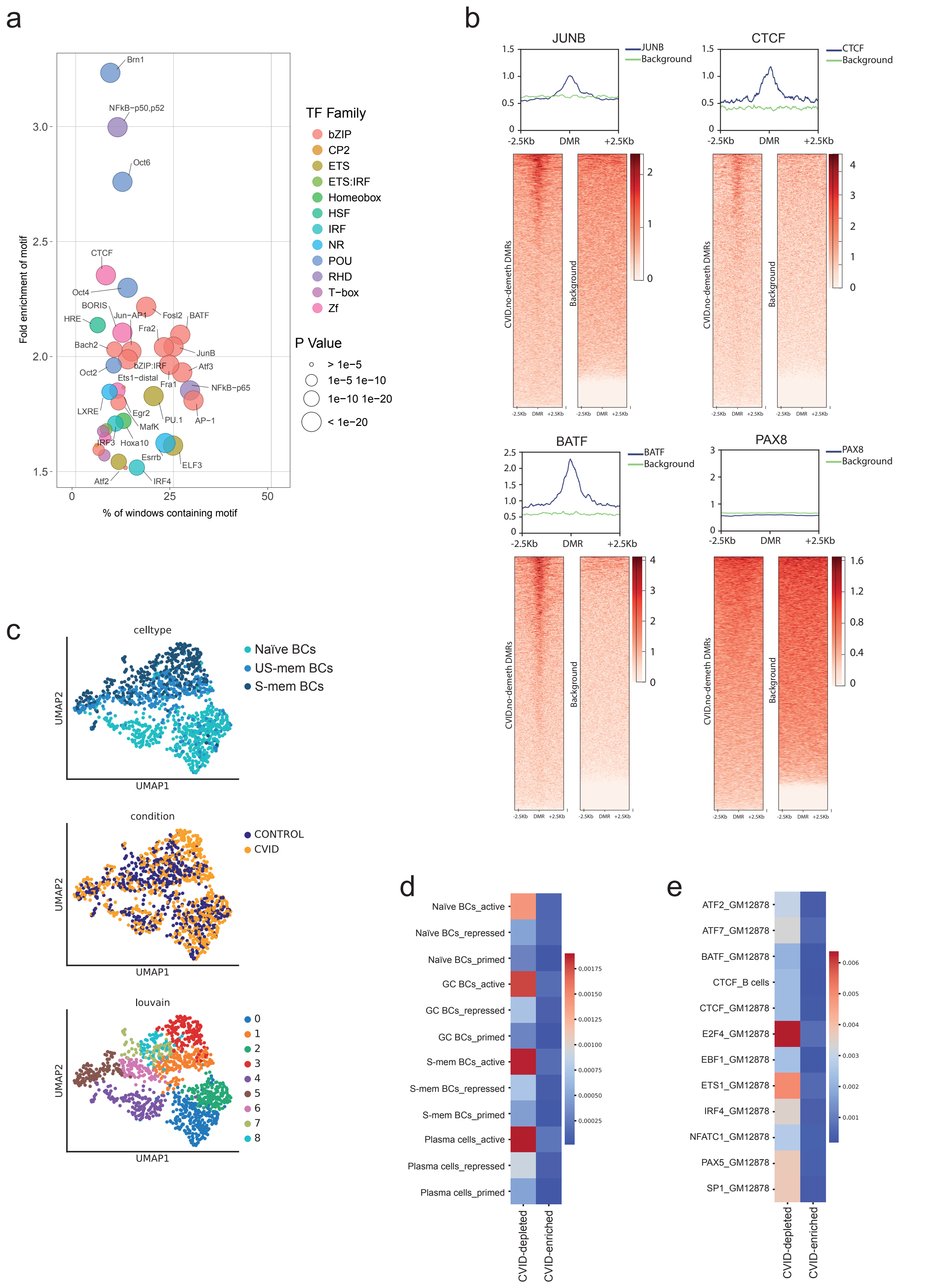

### Supp. Figure 5

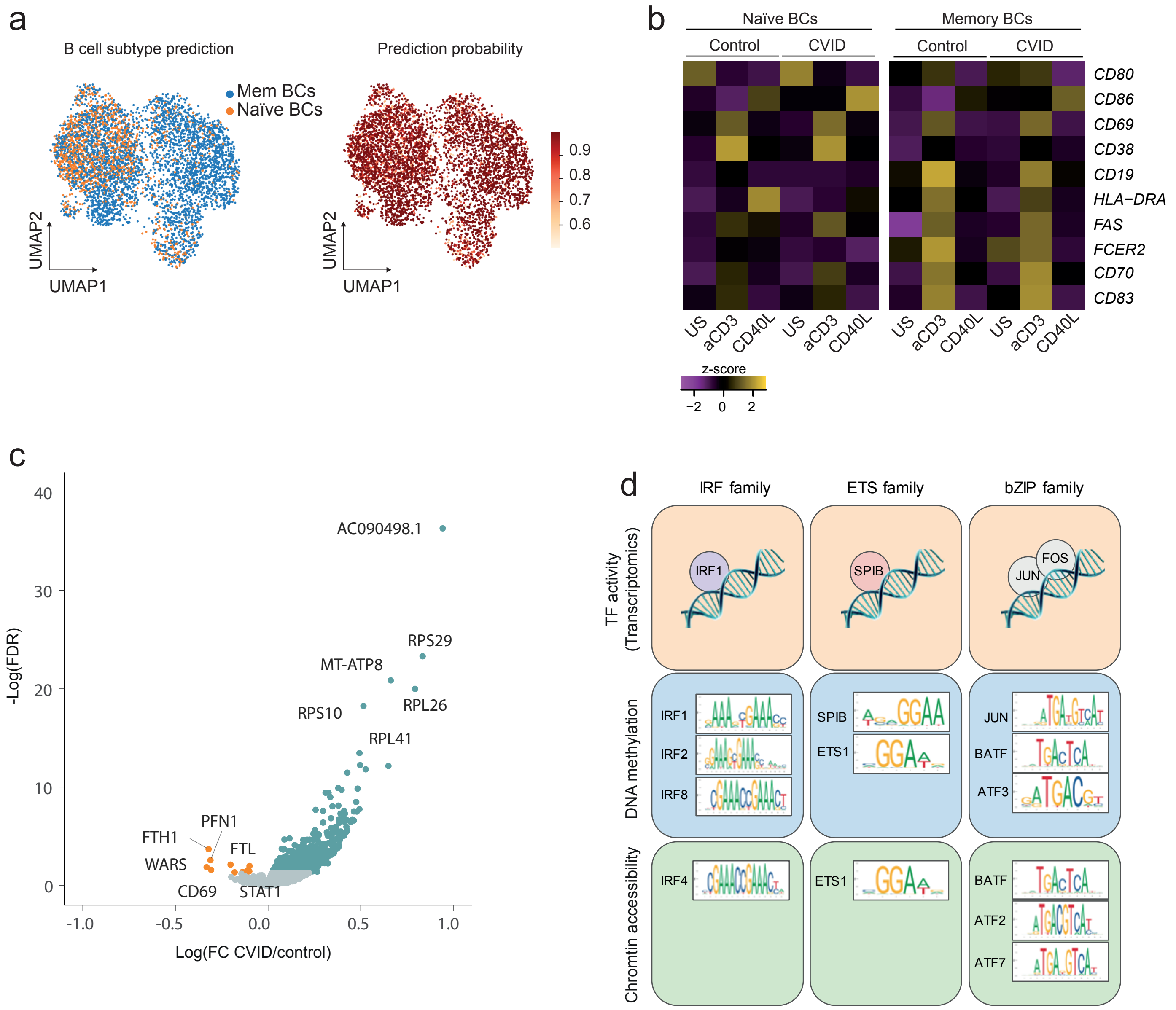

### Supp. Figure 6

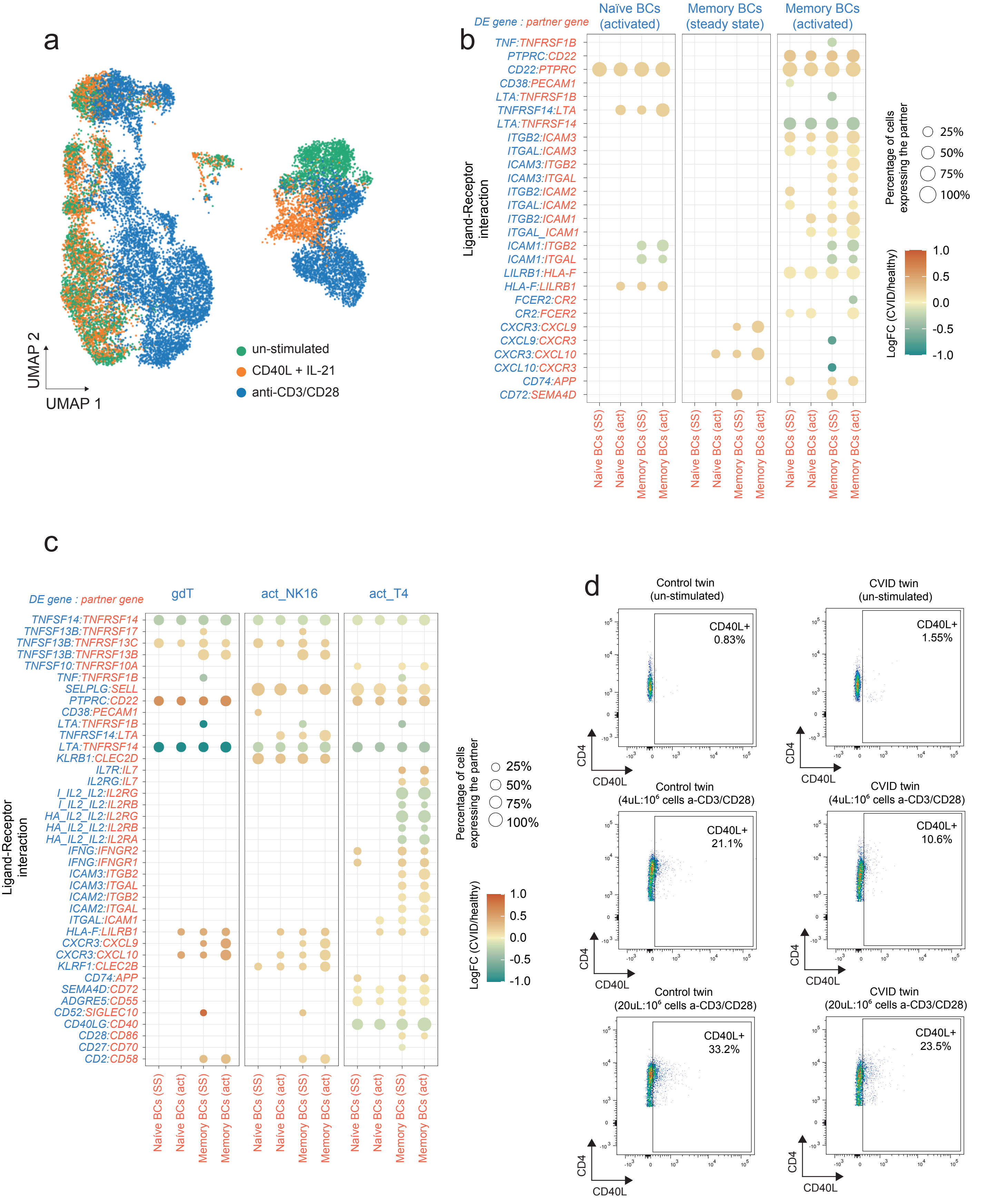

### Supp Figure 7

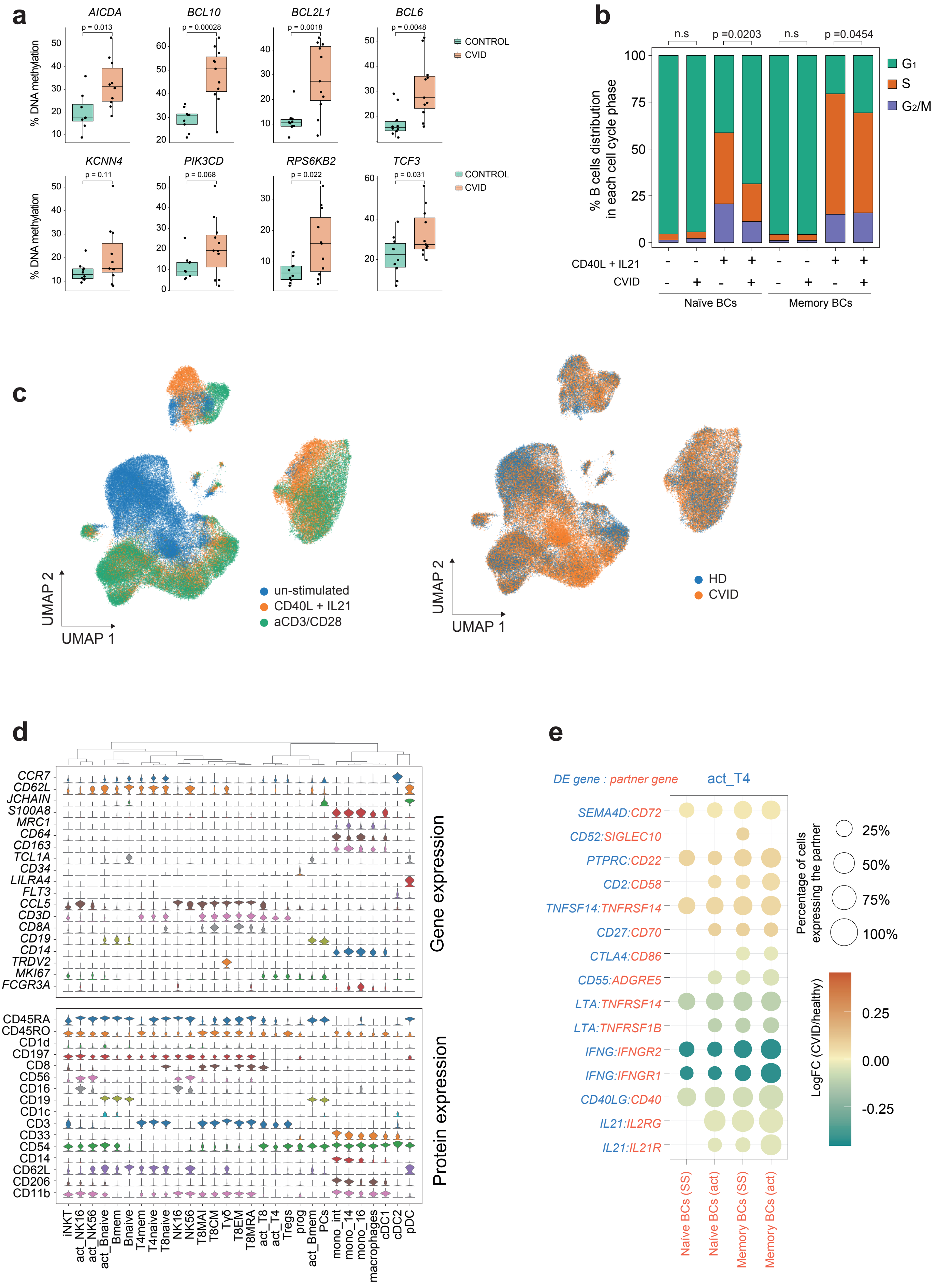

Supp. Figure 8

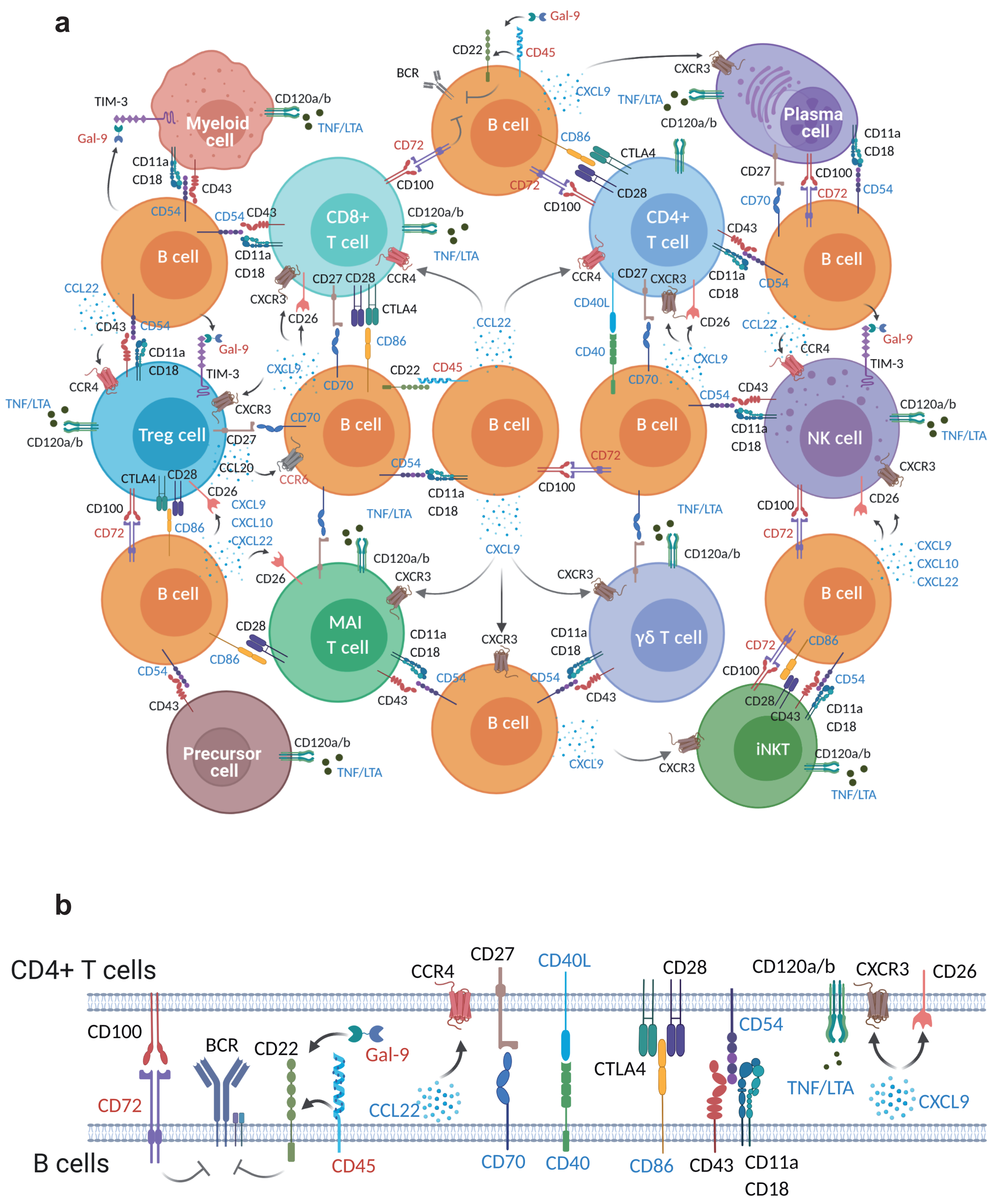

#### SUPPLEMENTARY FIGURE LEGENDS

**Supplementary Figure 1.** Pipeline of experiments and analysis performed. The techniques used are indicated inside green boxes, the analyses performed are indicated inside yellow boxes (with the softwares used in brackets), and the final output is indicated inside blue boxes.

**Supplementary Figure 2.** (a) FACS plots indicating the sorter strategy followed for naïve, US-mem and S-mem B cells isolation in control twin and (b) in CVID twin. (c) Absolute numbers and proportions of naïve, US-mem and S-mem B cells in the control and the CVID individual. The percentage of peripheral blood lymphocytes (PBL) and the percentage of total CD19+ B cells were represented.

**Supplementary Figure 3.** (a) Box plot representing global DNA methylation levels of the analyzed B cell subsets in 5 Mb windows throughout the genome (n=570 genomic regions analyzed on each B cell subset over one biologically independent sample from the control twin and one biologically independent sample from the CVID twin). Whiskers correspond with the minimum and maximum values of the data set (excluding any outliers). The box is drawn from Q1 to Q3 with a horizontal line to indicate the median. (b) Principal component analysis of DNA methylation in naïve (red), US-mem (blue) and S-mem (green) B cells from control and CVID twins. Dark colors indicate samples from the control twin and light colors indicate samples from the CVID twin. (c) Dot plots showing DNA methylation levels of windows containing 500 CpGs and Pearson correlation between control and CVID B cell compartments. (d) Bar plot showing the proportion of DMRs among the deciles of DNA replication time data from the immortalized B cell lines GM12878 and GM12801 (n=2 biologically independent samples). Bars represent the mean and error bars represent the SD. (e) Violin plots showing the expression of TET demethylating enzymes in naïve (n=174 biologically independent cells from the control twin and n=203 from the CVID twin), US-mem (n=227 biologically independent cells from the control twin and n=226 from the CVID twin) and S-mem (n=245 biologically independent cells from the control twin and n=249 from the CVID twin) B cells in the control and the CVID individual. No significant (n.s) differences were observed between both individuals using two-sided limma moderated t-test with Benjamini-Horchberg correction for multiple testing. (f) Bar plot depicting selected gene set enrichment analyzed using the Genomic Regions Enrichment of Annotations (GREAT) tool. The plot shows the most highly enriched terms associated with B cell biology from four annotation databases, based on probabilities from the binomial distribution. (g) Bar plot showing the genomic location of CVID.no-demeth DMRs.

**Supplementary Figure 4.** (a) Plot representing TF motif enrichment from CVID.no-demeth DMRs. Circle size indicates the significance; colors indicate the different TF families. (b) Heatmap and histogram representing the binding signal of several TFs (JUNB, CTCF and BATF) in human immortalized B cells at CVID.no-demeth DMRs ± 2.5 Kb flanking regions. PAX8 binding signal was used as a negative control. Transcription factors binding data were obtained from publicly available ChIP-seq datasets (see Methods). (c) UMAP visualization of unsupervised analysis of chromatin accessibility in naïve, US-mem and S-mem B cells from the control and CVID twin indicating the B cell subset, the origin of the cell and Louvain clustering. (d) Heatmap representing the association of the CVID-depleted and CVID-enriched DARs with different enhancer stages during B cell activation (active, primed and repressed enhancers). Active enhancer regions were defined for the concurrence of H3K4me1 and H3K27ac, repressed enhancers by H3K4me1 and H3K27me3, whereas primed enhancers by H3K4me1 exclusively. (e) Heatmap showing the association of CVID-depleted and CVID-enriched DARs with several TFs binding signal in human immortalized B cells and CTCF in human primary B cells.

**Supplementary Figure 5.** (a) Unsupervised UMAP visualizations of the CVID-discordant twins' B cell compartment indicating the prediction of the different B cells subsets and the probability of logistic regression prediction. (b) Heatmap showing the expression of different B cell activation genes in naïve and memory B cells from the control and the CVID twin. The gene expression in unstimulated cells (US) or cells tretated with anti-CD3/CD28 (aCD3) or CD40L + IL-21 (CD40L) is depicted. (c) Volcano plot representing differentially expressed genes in naïve B cells comparing control and CVID twins. The most differentially expressed genes are labeled. (d) Scheme depicting the common TFs implicated in the observed transcriptional, methylation, and chromatin accessibility alterations in the CVID-discordant twin pair.

**Supplementary Figure 6.** (a) UMAP plot showing the different immune cells analyzed and the corresponding cell stimulation protocol to which they were subjected. (b) Circle heatmaps representing selected dysregulated L/R interactions between B cells (autocrine/paracrine interactions) in the CVID twin. The scale indicates the $\log_2$(FC) gene expression of the different subsets of B cells in the CVID *vs.* control comparison. Only differentially expressed genes with an FDR < 0.05 were considered in the analysis. The percentage of B cells expressing the partner molecule is indicated by the circle size. Molecules of the L/R pairs expressed in one of the B cell subsets are shown in red; molecules of the L/R pairs expressed in the other B cells are shown in blue. Assays were carried out at the mRNA level, but were extrapolated to protein interactions. (c) Circle heatmaps representing selected dysregulated L/R interactions between B cells and the other immune cell compartments in the CVID twin. The scale indicates the $\log_2$(FC) gene expression of several non-B cells in the CVID *vs.* control comparison. Only differentially expressed genes with an FDR < 0.05 were considered in the analysis. The percentage of B cells expressing the partner molecule is indicated by the circle size. Molecules of the L/R pairs expressed in B cells are shown in red; molecules of the ligand-receptor pairs expressed in other non-B immune cells are shown in blue. Assays were carried out at the mRNA level, but were extrapolated to protein interactions. (d) FACS plots indicating the expression of CD40L in CD4+ T cells in unstimulated or stimulated PBMCs with anti-CD3/CD28 (4 µL/million cells or 20 µL/million cells). Due to the inherent limitations of these particular rare samples used for the FACS analysis, this is a single experiment which should not be taken as definitive proof.

**Supplementary Figure 7.** (a) Box plots showing the DNA methylation levels in S-mem B cells from the expanded cohort of CVID patients (n=11) and healthy controls (n=10) of several CpG sites/genes selected from our previous studies.[10,25] Whiskers correspond with the minimum and maximum values of the data set (excluding any outliers). The box is drawn from Q1 to Q3 with a horizontal line to indicate the median. A two-sided t-test was performed to detect significant differences in the DNA methylation levels between controls and CVID individuals. (b) Cell cycle distribution of naïve and memory B cells in CVID patients and healthy donors at steady state and stimulated with CD40L + IL-21, as estimated from single-cell datasets. One-sided Mann-Whitney test was performed to detect significant differences between the distribution of proliferating cells (S+G2M) between controls (n=8) and CVID individuals (n=10). (c) UMAP plot showing the different immune cells analyzed and the corresponding cell stimulation protocol to which they were subjected, as well as disease distribution. (d) Violin plots representing the expression of selected marker genes (top panel) and proteins (bottom panel) in the different cell clusters annotated. (e) Circle heatmaps representing selected dysregulated L/R interactions between B cells and activated CD4+ T cells in the CVID cohort. The scale indicates the $log_2$(FC) gene expression of activated CD4+ T cells in the CVID *vs.* healthy comparison. Only differentially expressed genes with an FDR < 0.05 were considered in the analysis. The percentage of B cells expressing the partner molecule is indicated by the circle size. Molecules of the L/R pairs expressed in B cells are shown in red; molecules of the ligand-receptor pairs expressed in activated CD4+ T cells are shown in blue. Assays were carried out at the mRNA level, but were extrapolated to protein interactions.

**Supplementary Figure 8.** (a) Scheme depicting the main dysregulated L/R interactions between B cells and other immune cell types in CVID. Names in blue correspond to downregulated genes in CVID and in red upregulated genes in CVID. (b) Scheme depicting the main dysregulated L/R interactions between B cells and the CD4+ T cell compartment in CVID. Names in blue correspond to downregulated genes in CVID and in red upregulated genes in CVID.
